# The 5-HT1A receptor antagonist WAY-100635 maleate reprograms metabolism to promote RGC differentiation and regeneration in retino-visual centers

**DOI:** 10.1101/2025.06.17.659983

**Authors:** Sayanta Dutta, Michelle L. Surma, Jie Chen, Kavitha Anbarasu, Jingwei Meng, Nian Wang, Arupratan Das

## Abstract

Metabolic collapse of retinal ganglion cells (RGCs) onsets glaucoma, yet no approved drug directly protects these neurons. Through a live-cell mitochondrial screen in human stem-cell-derived hRGCs we uncovered WAY-100635 (WAY), a clinically tested 5-HT1A antagonist, as a systemic neuroprotectant. WAY triggers a reversible cyclic-AMP surge that activates PGC-1α-driven reversible mitochondrial biogenesis and suppresses apoptosis. In glaucoma associated OPTN^E50K^ hRGCs, WAY restores mitochondrial fitness, dampens excitotoxicity, and reprograms metabolism toward aerobic glycolysis, while in progenitors WAY boosts mitochondrial cristae maturation, oxidative phosphorylation, and cell-cycle exit to accelerate RGC specification. Daily intraperitoneal dosing preserves RGC bodies, neural activity, promotes axon regeneration into the optic nerve and vision centers after optic-nerve crush, as well as shows RGC protection and maintenance of visual acuity in chronic ocular hypertension glaucoma. As the non-invasive neuroprotective therapy with a human safety profile, WAY addresses a critical gap in glaucoma care and potentially for other mitochondrial optic neuropathies.

**GRAPHICAL ABSTRACT:** 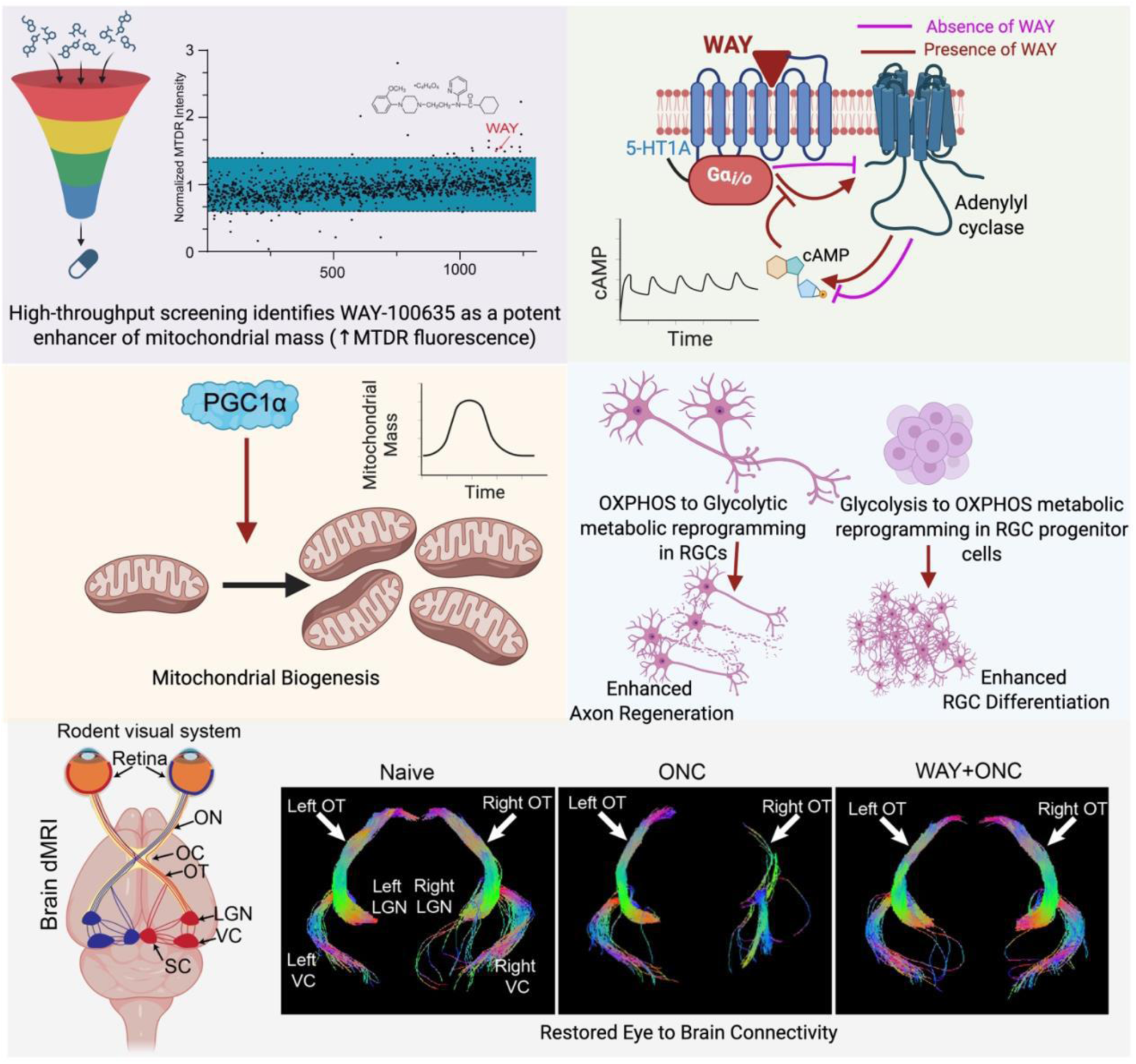

## INTRODUCTION

Neurons with long axons and partial unmyelination are particularly susceptible to mitochondrial dysfunctions. Retinal ganglion cells (RGCs) remain unmyelinated in the retina through the posterior end of lamina cribrosa which lack saltatory conduction and rely on densely packed mitochondria for efficient action potential propagation and transport, requiring high level of adenosine triphosphate (ATP)^1,2^. RGC degeneration stemming from mitochondrial abnormalities is a defining feature of both glaucoma and mitochondrial optic neuropathies (MON), including Leber hereditary optic neuropathy (LHON) and dominant optic atrophy (DOA)^2^.

While elevated intraocular pressure (IOP) has traditionally been regarded as the hallmark of glaucoma, approximately 40–50% of patients still experience progressive vision loss despite well-controlled IOP^3^. Moreover, more than half of primary open-angle glaucoma (POAG) patients present with RGC and vision loss in the context of normal IOP, often referred to as normal tension glaucoma (NTG)^4^. These clinical findings highlight the urgent need for therapeutic approaches that go beyond IOP management. Glaucoma is axogenic in nature, initiated by distal axonal degeneration in the brain with gradual progression towards the proximal end and eventual cell body death. Hence, early-stage interventions to protect or restore axonal function represent a promising avenue for halting disease progression^5^. This has been shown in glaucoma models where increased IOP disrupts mitochondrial function^6^ and depletes ATP^7^, impairing axonal transport and ultimately leading to RGC death^5,8,9^.

Mitochondrial dysfunction is increasingly recognized as a fulcrum of RGC degeneration, making mitochondrial repair an attractive therapeutic entry point. Genetic strategies that enforce mitochondrial fusion^10^ or boost anterograde transport by over-expressing Armcx1^11^, Disc1^12^, or the Optineurin (OPTN) associated TRAK1-KIF5B motor complex^13^, drive impressive RGC axon regrowth in the optic nerve after optic-nerve crush (ONC). Likewise, metabolic augmentation with pyruvate^14^, nicotinamide/Vitamin B₃, NAD⁺, or Nmnat1 gene therapy^15,16^ attenuates RGC loss in models of elevated intra-ocular pressure (IOP). However, chronic mitochondrial fusion or relentless distal transport can disrupt healthy dynamics and deplete mitochondria in the proximal axonal regions, while sustained oxidative phosphorylation (OXPHOS) by mitochondria risks toxic oxidative by-products, limiting the long-term benefit. Most importantly, no current intervention has restored functional eye-to-brain connectivity, which is a prerequisite for glaucoma therapy. A non-invasive, metabolism-restorative small molecule capable of rebuilding the retino-visual pathway would therefore constitute a transformative leap toward durable neuroprotection in glaucoma.

Here, we combined phenotypic screening with human stem-cell–derived retinal ganglion cells (hRGCs) to pinpoint small molecules that restore mitochondrial fitness. The screen revealed WAY-100635 maleate (WAY), a well-studied 5-HT1A Gi/o-coupled receptor antagonist^17,18^, as a potent, reversible inducer of cyclic AMP and PGC-1α–driven mitochondrial biogenesis. WAY confers metabolic reprogramming that accelerates hRGC differentiation and grants strong neuroprotection to hRGCs harboring the glaucoma-linked OPTN^E50K^ mutation^19^. The OPTN^E50K^ mutation is found among a severe form of NTG patients characterized by defective mitochondrial homeostasis^19,20^. In vivo, a systemic course of WAY preserves RGC somata, drives axon regrowth through the optic nerve and central visual targets after crush injury, and safeguards both RGC survival and visual acuity in glaucoma models. Crucially, WAY has already cleared FDA toxicology hurdles as a serotonin-receptor probe in clinical trial (NCT00603018, NCT02810717), streamlining its path toward clinical deployment.

Collectively, our data establish a first-in-class pharmacologic blueprint for simultaneously protecting and regenerating RGC axons by modulating mitochondrial dynamics, achieved without the constitutive changes or delivery barriers of gene therapy. Because mitochondrial failure is a shared lesion across neurodegenerative disorders, this strategy may extend to Parkinson’s disease, amyotrophic lateral sclerosis, and beyond, expediting translation through repurposing of a drug with an established safety record.

## RESULTS

### Identification of WAY-100635 that activates neuroprotective signaling in hRGCs through antagonizing 5-HT1A receptor

Building on our recent discovery that boosting mitochondrial biogenesis safeguards human stem-cell derived hRGCs, including those carrying the glaucoma-linked OPTN^E50K^ mutation^20^, we performed a high-throughput screen for druggable enhancers of mitochondrial health. We screened the LOPAC (Library of Pharmacologically Active Compound, Sigma) library in highly purified (>90 %) hRGC cultures that recapitulate native RGC transcriptional and electrophysiological signatures, a method widely accepted in the field, and we routinely use^20–22^. Mitochondrial fitness was quantified by flow-cytometric measurement of MitoTracker Deep Red (MTDR) fluorescence, a sensitive readout of membrane-potential– positive mitochondrial mass^23^. Compounds that elevated single-cell MTDR signal 40% above the median threshold were flagged as primary hits (**Fig. 1a**). Secondary triage required that a hit both (i) maintain an elevated MitoTracker Deep-Red signal and (ii) lower the apoptosis (cleaved-caspase-3 activity) in hRGCs. Only a few compounds met both benchmarks, and WAY-100635 maleate (WAY) delivered the most pronounced dual response (**Fig. 1b, c**). Coupled with its documented safety in human studies, these data positioned WAY as the lead candidate for downstream glaucoma-neuroprotection analyses. Notably, under culture conditions even wild-type (Wt) hRGCs suffer apoptosis over time^24^; WAY significantly reduced this baseline apoptosis, indicating a broad neuroprotective action independent of any obvious genetic defects which is often the case for POAG. WAY is a well-characterized antagonist of the Gi/o-coupled 5-HT1A receptor. To test receptor specificity, we treated hRGCs with the bona fide 5-HT1A agonist 8-OH-DPAT (DPAT)^25,26^. DPAT failed to diminish apoptosis (**Fig. 1d**), underscoring that the protective effect is linked to 5-HT1A antagonism by WAY rather than generic serotonergic signaling. Together, these findings warrant WAY as a potent, mitochondria-restorative small molecule and establish 5-HT1A antagonism as a tractable axis for RGC neuroprotection.

**Fig. 1.**
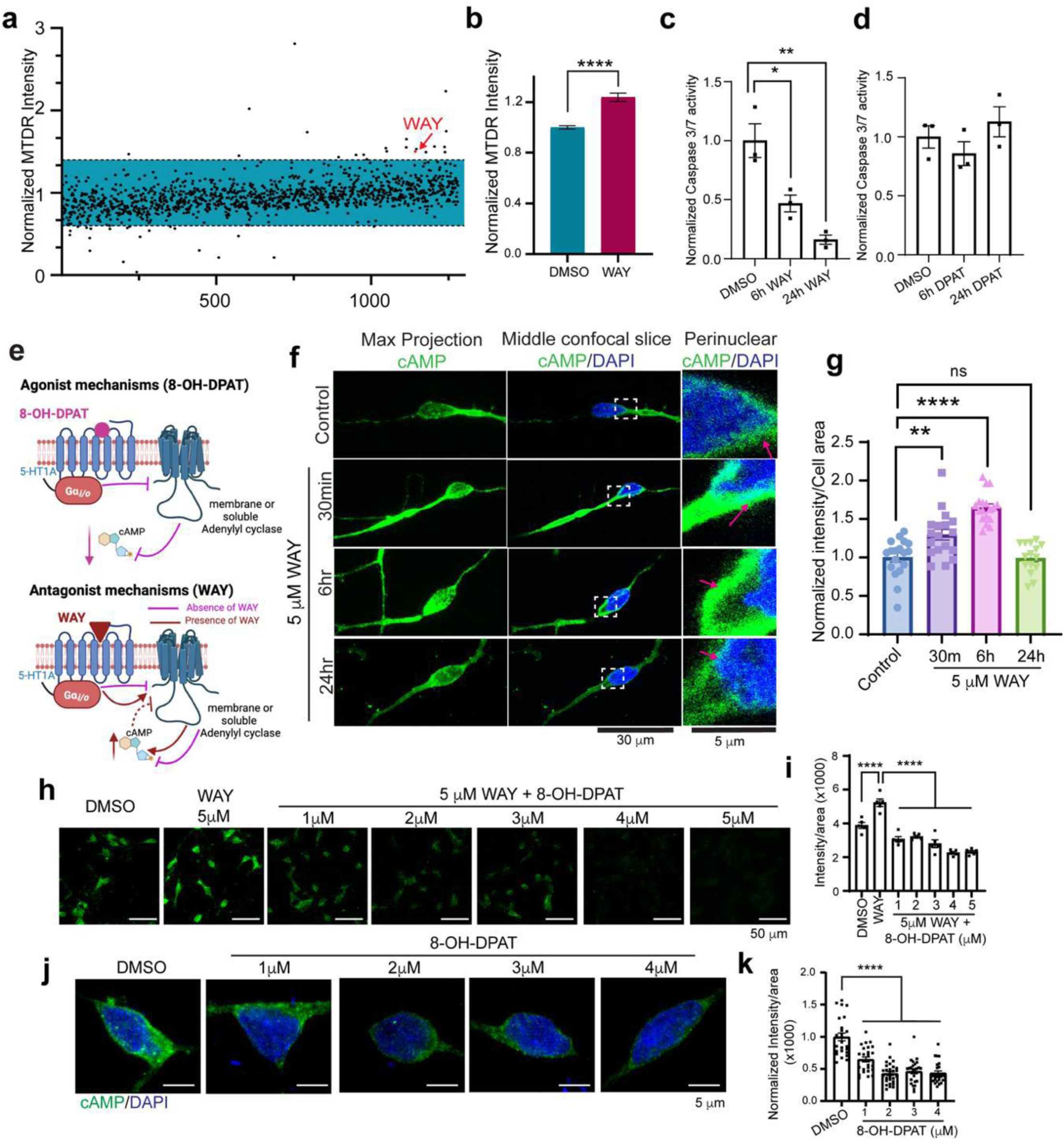
WAY-100635 engages 5-HT1A receptor to reversibly stimulate neuroprotective second messenger cAMP. (**a**) High-content screen of H7-hRGC^WT^ (derived from the H7-hESC reporter line) plotted fold-change in MitoTracker Deep Red (MTDR) fluorescence; compounds altering mitochondrial mass by more than 40 % lie outside the blue box, with WAY-100635 (WAY) highlighted in red. (**b**) Independent validation (5 µM, 24 h) confirms increased MTDR signal relative to untreated control (CTR) (unpaired t-test, **** p < 0.0001; n = 3 biological replicates; mean ± SEM). (**c, d**) Caspase-3/7 activity (ApoToxGlo-Triplex) after 24 h treatment with WAY (5 µM) or the 5-HT1A agonist 8-OH-DPAT (DPAT, 5 µM), expressed as normalized to CTR (one-way ANOVA, Dunnett’s post-hoc; * p < 0.05, ** p < 0.01, *** p < 0.001, **** p < 0.0001; n = 3). (**e**) Cartoon summarizing reciprocal cAMP signaling elicited by 5-HT1A antagonism (WAY) or agonism (DPAT). (**f, g**) Confocal immunofluorescence with anti-cAMP antibody (max projection and central z-plane) shows time-dependent perinuclear cAMP accumulation after WAY. Quantification of cAMP intensity per cell area from sum projections, normalized to control from 15–18 cells per condition (one-way ANOVA, Dunnett’s). (**h, i**) WAY pre-treatment (1 h, 5 µM) elevates cAMP, which is dose-dependently suppressed by subsequent DPAT (1 -5 µM, 6 h). Analysis from 5 images per condition (∼ 80-150 cells per image; one-way ANOVA, Tukey post-hoc). (**j, k**) DPAT alone (6 h) lowers basal cAMP (25-29 cells per condition; one-way ANOVA, Dunnett’s post-hoc). Quantification done from sum projections as in **g**. Error bars represent SEM throughout. DMSO served as vehicle for DPAT.

RGCs endogenously release serotonin, a 5-HT1A agonist that engages Gi/o proteins, suppresses its cognate adenylyl-cyclase (AC) activity, and enforces low intracellular cyclic adenosine mono phosphate (cAMP) levels^27,28^. Accordingly, hRGCs exposed to serotonin, or the canonical agonist DPAT, this is expected to display diminished cAMP, whereas the antagonist WAY is predicted to competetively inhibit agonist binding leading to disinhibition of AC and cAMP elevation (**Fig. 1e**). Consistent with this model, WAY treatment elicited a rapid but fully reversible rise in cAMP that decayed to baseline with classical GPCR kinetics (**Fig. 1f, g**). Strikingly, the elevation was enriched in the perinuclear compartment (**Fig. 1f**), a spatial signature previously linked to neuroprotection^29,30^. Because these experiments were performed in highly purified hRGC cultures devoid of other cell types, the response is unequivocally cell-autonomous. Continuous exposure to WAY for six days produced repeated, non-desensitizing oscillations in cAMP (**Fig. S1**), demonstrating that 5-HT1A antagonism can engage downstream signaling without driving irreversible, constitutive changes, an essential property for therapeutic use.

To confirm receptor specificity, we performed a competitive reaction to 5-HT1A by the agonist DPAT and antagonist WAY. Competitive binding studies in primate brain have established that both WAY-100635 (WAY) and DPAT engage 5-HT1A with high selectivity^31^. In hRGCs, WAY alone elevated cAMP, whereas DPAT alone lowered it, responses predicted for antagonism versus agonism at a Gi/o-coupled receptor (**Fig. 1f, g, h-k**). Co-application of DPAT completely abolished the WAY-induced cAMP rise (**Fig. 1h, i**), indicating direct competition at the same binding site. Although limited reports suggest off-target activity of WAY at DRD4 (Gi/o-coupled)^32,33^ and DPAT at 5-HT7 (Gs-coupled)^34,35^ as agonist, such interactions in hRGCs would have produced the opposite cAMP signatures, decrease for WAY and increase for DPAT, which we did not observe. These data demonstrate that WAY’s neuroprotective action in hRGCs is mediated specifically through 5-HT1A antagonism.

### WAY restores mitochondrial homeostasis to protect hRGCs

The increase in MTDR fluorescence observed after WAY exposure could reflect either enhanced mitochondrial biogenesis or a boost in membrane potential within the existing network. To distinguish these possibilities, we quantified mitochondrial mass during a WAY treatment time course by (i) confocal immunofluorescence using antibodies against the outer-membrane protein Tom20 and (ii) immunoblotting for Tom70. The imaging assay revealed a reversible surge in mitochondrial content that peaked at 6 h and returned to baseline by 24 h (**Fig. 2a–b**). The imaging peak coincided with maximal cAMP elevation, whereas the immunoblot assay showed reversible surge, but the peak lagged by ∼12 h (**Fig. 2c–d**), presumably due to the lag in protein turnover kinetics. Because nuclear PGC-1α orchestrates mitochondrial biogenesis and can be activated by cAMP^36^, we examined its subcellular distribution. Confocal analyses showed a parallel, reversible enrichment of PGC-1α in nuclei (**Fig. 2e, f**), with kinetics that mirrored the 6-h biogenesis pulse. Continuous replenishment of WAY for 6 days in the medium maintained this cyclic response for cAMP (**Fig. S1**), indicating that the transient profile is intrinsic to the signaling cascade rather than drug depletion. To test whether PGC-1α is required for the mitochondrial biogenic pulse, we depleted PGC-1α with antisense oligonucleotides (ASOs) targeting the 3′ UTR (**Fig. 2g**). Control ASO-treated cells retained the characteristic rise-and-fall in mitochondrial mass after WAY (**Fig. 2h, i**). Strikingly, PGC-1α knockdown abolished this effect (**Fig. 2j, k**) and concomitantly blunted WAY-mediated protection against apoptosis (**Fig. 2l**). Collectively, these data demonstrate that WAY elicits a short-lived, cAMP-driven burst of mitochondrial biogenesis that is PGC-1α dependent and functionally linked to the compound’s neuroprotective action in hRGCs.

**Fig. 2.**
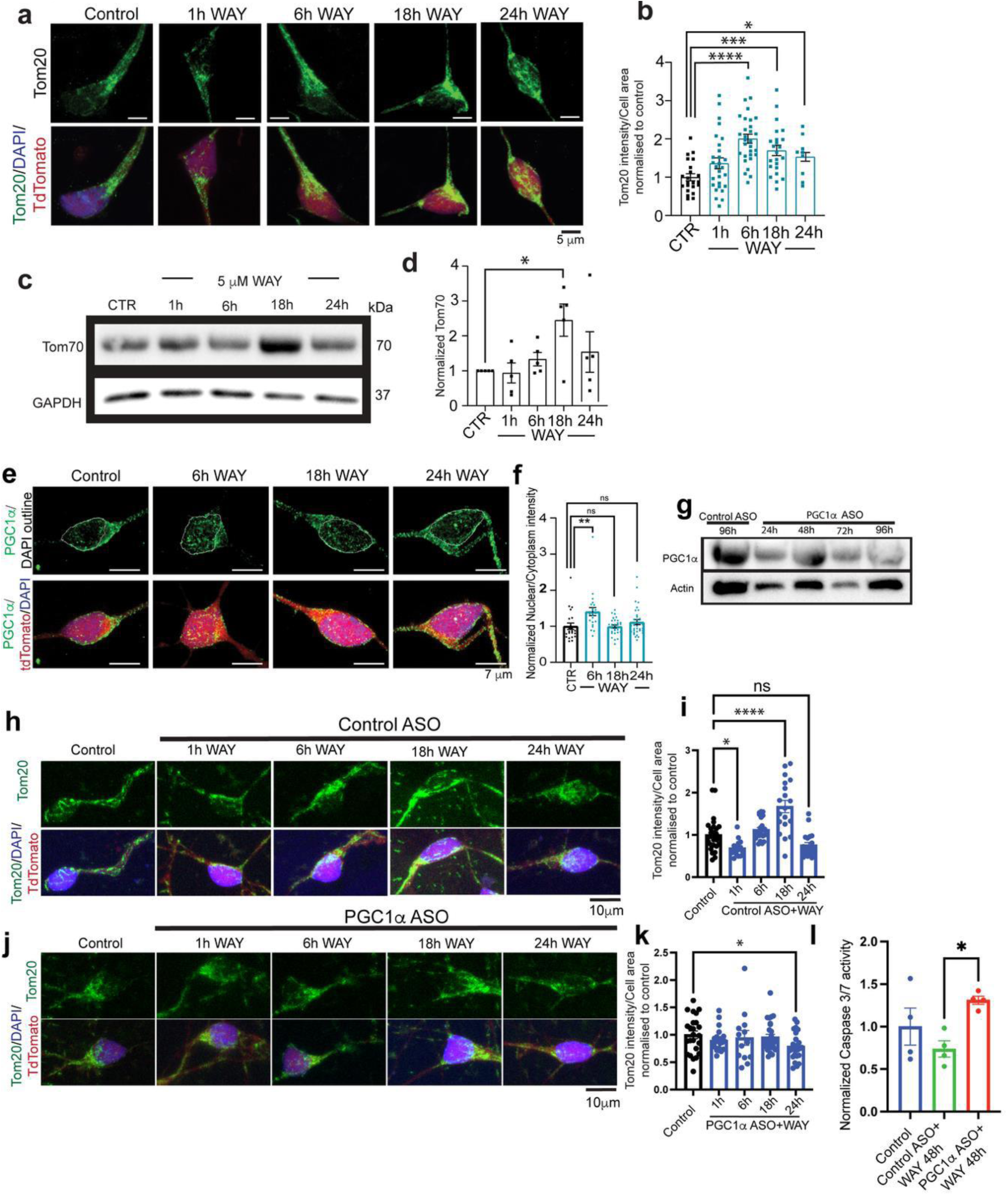
WAY-100635 reversibly induces mitochondrial biogenesis in hRGCs via PGC-1α. (**a**) Representative confocal images (max projection of z-stacks) of H7-hRGC^WT^ expressing tdTomato (red), and stained for nucleus (DAPI, blue), mitochondria (Tom20, green) at the indicated times after 5 µM WAY. (**b**) Tom20 fluorescence per cell area (sum projection) expressed as normalized to untreated control (CTR). One-way ANOVA, Dunnett post-hoc; * p < 0.05, *** p < 0.001, **** p < 0.0001; n = 19 - 30 cells; (**c, d**) Immunoblot and densitometry for Tom70 after WAY (5 µM). Values are Tom70/GAPDH, normalized to CTR (n = 5; one-way ANOVA, Dunnett post-hoc * p < 0.05). (**e, f**) Confocal images (max projection) and quantification of PGC-1α nuclear translocation following WAY (5 µM). Data are nuclear-to-cytoplasm intensity ratios from sum projections normalized to CTR (n = 22–33 cells; one-way ANOVA, Tukey post-hoc; * p < 0.05, ** p < 0.01, *** p < 0.001, **** p < 0.0001). (**g**) Time-course immunoblot validating knock-down of PGC-1α with a 3′-UTR-targeting antisense oligonucleotide (ASO) versus non-targeting control. (**h - k**) Confocal image (max projection) of H7-hRGC^WT^ stained for mitochondria (Tom20) after 24 h control or PGC-1α ASO pretreatment followed by WAY (5 µM, 1 -24 h). Control ASO permits the expected rise in mitochondrial mass (**h, i**); PGC-1α ASO blocks it (**j, k**). Quantification of Tom20 intensity done per cell area from sum projections (n = 13 - 29 cells; one-way ANOVA, Dunnett’s post-hoc; * p < 0.05, **** p < 0.0001). (**l**) Caspase-3/7 activity (ApoToxGlo-Triplex) after 24 h ASO pretreatment + 48 h WAY, plotted as normalized to CTR (n = 4; one-way ANOVA, Tukey; * p < 0.05). Error bars represent SEM.

WAY reversibly activated mitochondrial biogenesis in WT hRGCs. We therefore asked whether the same response occurs in NTG-associated OPTN^E50K^ hRGCs. Confocal immunofluorescence for the mitochondrial marker Tom20 revealed a transient rise in mitochondrial mass that peaked at 6 h, identical to the kinetics observed in WT cells (**Fig. 3a, b**). We showed that OPTN^E50K^ hRGCs harbor fewer, metabolically over-burdened mitochondria, but stimulating biogenesis via TBK1 activation relieves ATP demand and reduces apoptosis^20^. Consistent with this model, WAY increased mitochondrial membrane potential, measured with the potentiometric dyes JC-1 (red/green ratio) (**Fig. 3c, d**), and TMRM (mitochondrial area per cell), across different treatment time points (**Fig. 3e, f**).

**Fig. 3.**
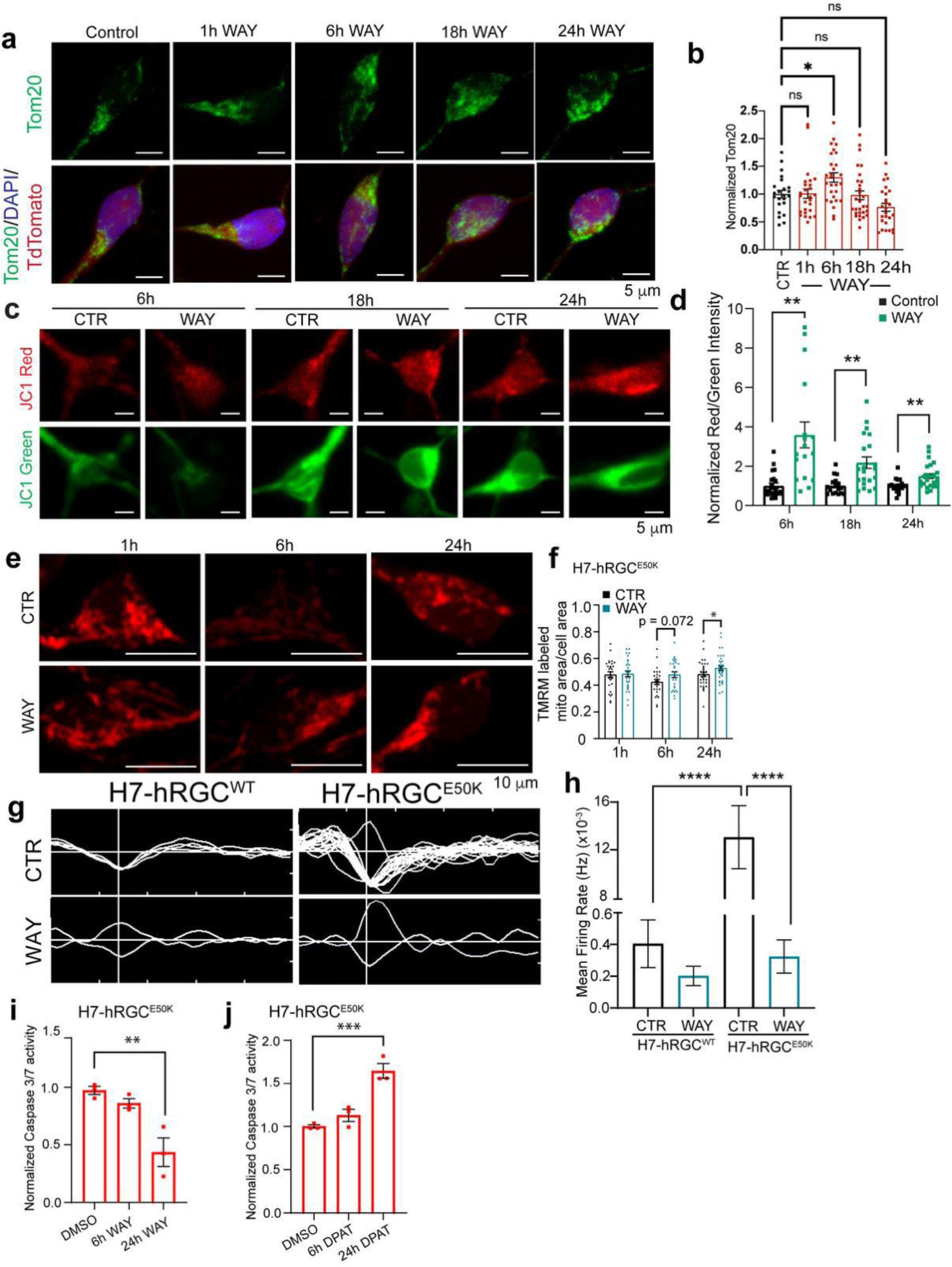
**WAY-100635 restores mitochondrial homeostasis to provide protection to glaucomatous OPTN^E50K^ hRGCs**. (**a**) Representative confocal images (max projection of z-stacks) of H7-hRGC^E50K^ expressing tdTomato (red), and stained for nucleus (DAPI, blue), mitochondria (Tom20, green) at the indicated times after 5 µM WAY treatment. (**b**) Tom20 fluorescence per cell area from sum projections, expressed normalized to untreated control (CTR). One-way ANOVA, Dunnett post-hoc; * p < 0.05; n = 23 - 29 cells. (**c, d**) Mitochondrial membrane potential assessed with JC-1. Live-cell images (**c**) and red/green fluorescence ratios normalized to corresponding control ratios (**d**) show a significant increase after WAY (Student’s t-test; ** p < 0.01, *** p < 0.001, **** p < 0.0001; n = 17–24). (**e, f**) Mitochondrial membrane potential confirmation with TMRM labeling: representative live cell images (**e**) and quantified fluorescence per cell area from sum projections (**f**) (Student’s t-test; * p < 0.05, **** p < 0.0001; n = 25–35). (**g**) Representative action potential firing spikes from a single micro-electrode array (MEA) sensor over 3 min under the indicated conditions. (**h**) Mean firing rate (16 sensors/well) pooled across wells (Two-way ANOVA, Tukey post-hoc; **** p < 0.0001; n = 4–5 wells). (**i, j**) Caspase-3/7 activity (ApoToxGlo-Triplex) after 48 h treatment with WAY (5 µM) or DPAT (5 µM), normalized to CTR (one-way ANOVA, Dunnett’s post-hoc; ** p < 0.01, *** p < 0.001; n = 3).

Mitochondrial depolarization drives excitotoxic hyperactivity^37^. Multi-electrode array recordings confirmed that OPTN^E50K^ hRGCs fire at higher spontaneous rates than WT controls, echoing prior patch-clamp studies^38^; remarkably, WAY normalized firing frequency to WT levels (**Fig. 3g, h**). Corroborating its neuroprotective effect, WAY reduced apoptosis in OPTN^E50K^ hRGCs, whereas the 5-HT1A agonist DPAT increased apoptosis (**Fig. 3i, j**). The OPTN gene is critical for mitochondrial homeostasis, and its E50K allele appears in ∼17 % of NTG patients with aggressive disease^19^, highlighting its therapeutic relevance.

Together, these data show that antagonism of 5-HT1A by WAY transiently boosts mitochondrial biogenesis, restores mitochondrial health, suppresses excitotoxic activity, and ultimately protects OPTN^E50K^ hRGCs, highlighting a potential therapeutic avenue for normal-tension glaucoma.

### WAY orchestrates cell-state specific metabolic reprogramming driving neuroprotective glycolysis in mature RGCs while enhancing oxidative phosphorylation in progenitor stem cells to accelerate differentiation

Under normal conditions, RGCs derive most of their ATP from mitochondrial oxidative phosphorylation (OXPHOS), however, excessive oxidative stress associated with OXPHOS promotes RGC degeneration^39^. Because WAY reversibly enhanced mitochondrial biogenesis and lowered apoptosis in both WT and OPTN^E50K^ hRGCs, we asked whether these benefits involve metabolic reprogramming. In the Seahorse Mito Stress Tests, that measures mitochondrial respiration, we observed higher oxygen consumption rate (OCR) for OPTN^E50K^ hRGCs for both control and WAY treatment under basal and FCCP induced maximum mitochondrial respiration (**Fig. 4a**). This led to the higher basal respiration (**Fig. 4b**) and mitochondrial ATP (MitoATP) synthesis for the OPTN^E50K^ hRGCs than WT cells but without having additional effects of WAY treatment (**Fig. 4c**). Increased mitochondrial ATP production for OPTN^E50K^ hRGCs is consistent to our earlier findings^20^, but this is intriguing that even though WAY reversibly induces mitochondrial biogenesis and restores mitochondrial homeostasis it does not promote more mitochondrial respiration. This can be due to metabolic reprogramming in which WAY rebalances metabolic load between mitochondria and glycolysis to limit OXPHOS related oxidative stress, yet meeting the ATP requirements for cellular physiology. To obtain direct evidence for change in glycolysis rate, we conducted Glycolytic-Rate Assay. In this assay seahorse analyzer measures extracellular acidification rate (ECAR) originates from cellular proton efflux and OCR to measure OXPHOS. Extracellular protons originate from glucose metabolism to lactate by glycolysis and by hydration of CO_2_ that produced by mitochondrial respiration. Inhibition of mitochondrial electron transport chain by Rot/AA inhibits proton source from mitochondria but drives maximum glycolytic capacity hence corresponding proton efflux rate (PER) as compensatory ATP source. Next, the addition of 2-deoxy-D-glucose (2-DG) a glucose analog that inhibits glycolysis, drops PER confirms the proton efflux prior to 2-DG is from glycolysis (**Fig. 4d**). The fraction of proton efflux rate (PER) attributable to glycolysis remained similar in WT cells and significantly increased in the OPTN^E50K^ hRGCs under WAY treatment (**Fig. 4d, e**). Accordingly, the mitoOCR/glycoPER ratio declined moderately in WT and markedly in the OPTN^E50K^ hRGCs (**Fig. 4f**) reflecting glycolytic metabolic state. Following 2-DG application, residual acidification is not attributable to mitochondria but any residual glycolysis that are not inhibited. Post-2-DG acidification rose significantly in OPTN^E50K^ hRGCs without any effect on WT cells, implying higher basal glycolysis that is not completely inhibited (**Fig. 4g**). Collectively, these data show that WAY restores mitochondrial health yet paradoxically redirects cellular energetics toward aerobic glycolysis, avoiding further OXPHOS stimulation in the glaucoma associated OPTN^E50K^ hRGCs. The pronounced shift toward aerobic glycolysis we uncover here mirrors protective metabolic adaptations reported across diverse neurodegenerative models including optic nerve injury^40,41^.

**Fig. 4.**
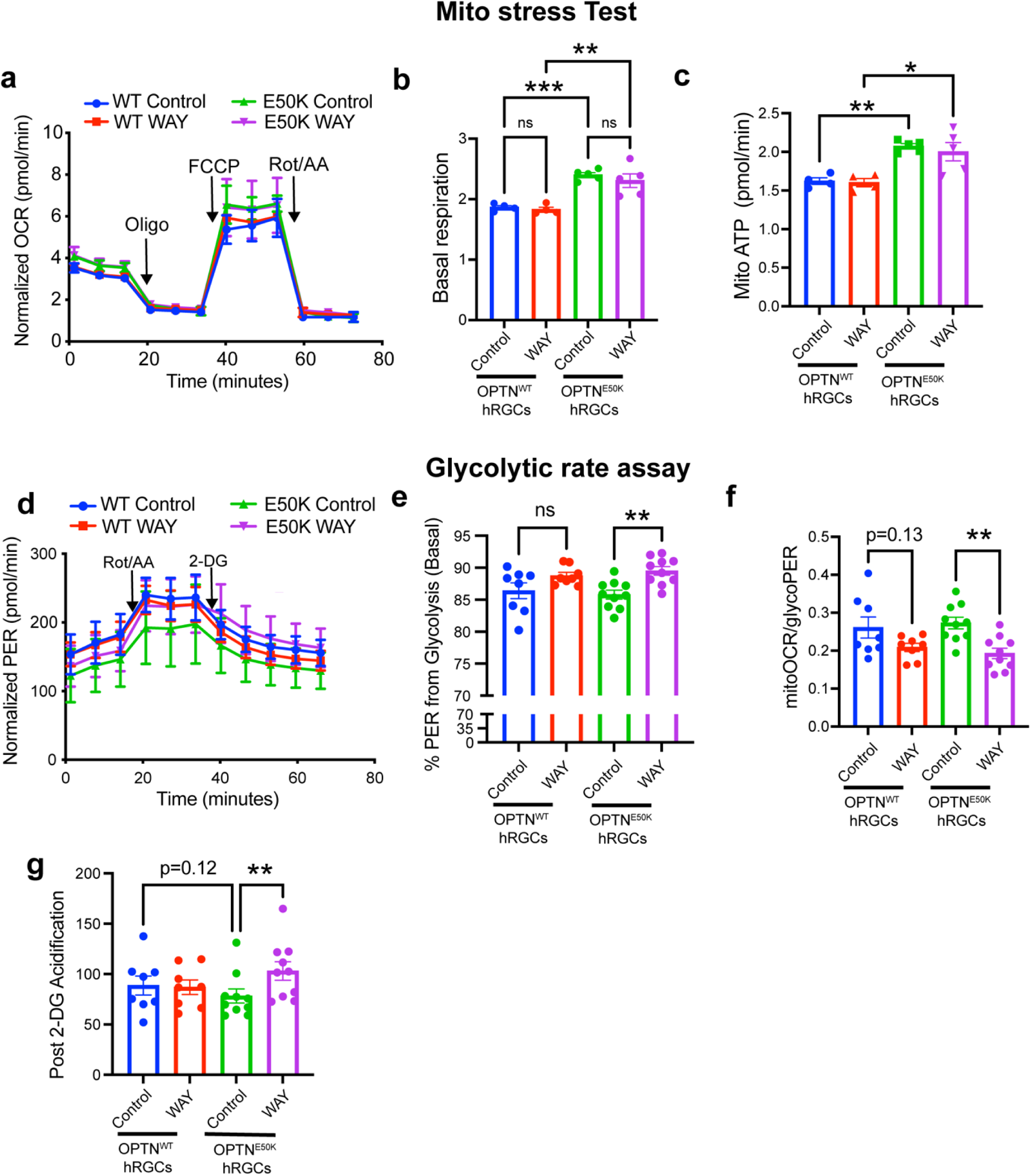
WAY-100635 induces glycolytic metabolic reprograming in hRGCs. **(a**) Cell area normalized oxygen consumption rate (OCR) measured by Mito Stress Test in H7-hRGC^WT^ and H7-hRGC^E50K^ RGCs treated with or without WAY-100635 for 24 h. (**b - c**) Quantification of basal respiration and mitochondrial ATP production (pmol/min) in respective RGCs. Two-way ANOVA with Tukey; * p < 0.05, ** p < 0.01, *** p < 0.001; n = 4-5 biological repeats. (**d**) Cell area normalized proton efflux rate (PER) from the Glycolytic Rate Assay in H7-hRGC^WT^ and H7-hRGC^E50K^ RGCs cells treated with or without WAY for 24 h. (**e - g**) Quantification of basal percentage of PER from glycolysis (**e**), mitoOCR/glycoPER ratio (**f**), and post 2-DG acidification (**g**) in respective RGCs. Two-way ANOVA with Tukey; ** p < 0.01; n = 8-10 biological repeats; Error bars are SEM.

Cues that drive pluripotent progenitors toward a neuronal fate can, in parallel, endow nascent neurons with heightened stress resistance^42^. Because stem cells maintain a low immature mitochondrial network, predominantly relying on glycolysis for ATP source^43^, we asked whether stimulating mitochondrial biogenesis at this stage would enhance hRGC differentiation. We observed, hESCs treated with WAY-100635 (5 µM) displayed a robust increase in mitochondrial content, quantified by Tom20 immunofluorescence and Tom70 immunoblotting (**Fig. S2a-d**). In contrast to the reversible rise observed in differentiated hRGCs, mitochondrial mass in hESCs progressively elevated during the 24-h time course, indicating a lineage-specific response to WAY. We next interrogated the developmental window during which WAY may enhance differentiation. The small molecule-based differentiation using RGC reporter stem cells used here produces tdTomato-positive hRGCs at around day 30 and reaches maximum number around day 45^22^. Cultures were pulsed with WAY (2.5 µM or 5 µM) for 4-day intervals between days 2 and 26, and tdTomato-positive hRGCs were quantified on day 32 using our BRN3B-P2A-tdTomato-P2A-Thy1.2 reporter line. A single exposure during days 2–6 produced ∼1.5-fold increase in tdTomato⁺ cells at both concentrations, with a modest but significant effect for days 6–10 at 2.5 µM; later windows were ineffective as revealed by imaging (**Fig. S3a**) and by quantitative flow cytometry analysis (**Fig. S3b, c**). Expanding the dose range (1 - 2.5 µM) within the days 2-6 window reproducibly elevated hRGC output irrespective of concentration, as shown by microscopy (**Fig. 5a**) and quantitative flow cytometry analysis (**Fig. 5b, c**). Molecular markers independent of tdTomato positive readouts further corroborated enhanced differentiation as early WAY treatment up-regulated BRN3B and SIX3 transcripts, key determinants of RGC lineage commitment^44,45^ and ocular neuroectoderm patterning^46,47^, respectively relative to vehicle controls (**Fig. 5d**).

**Fig. 5.**
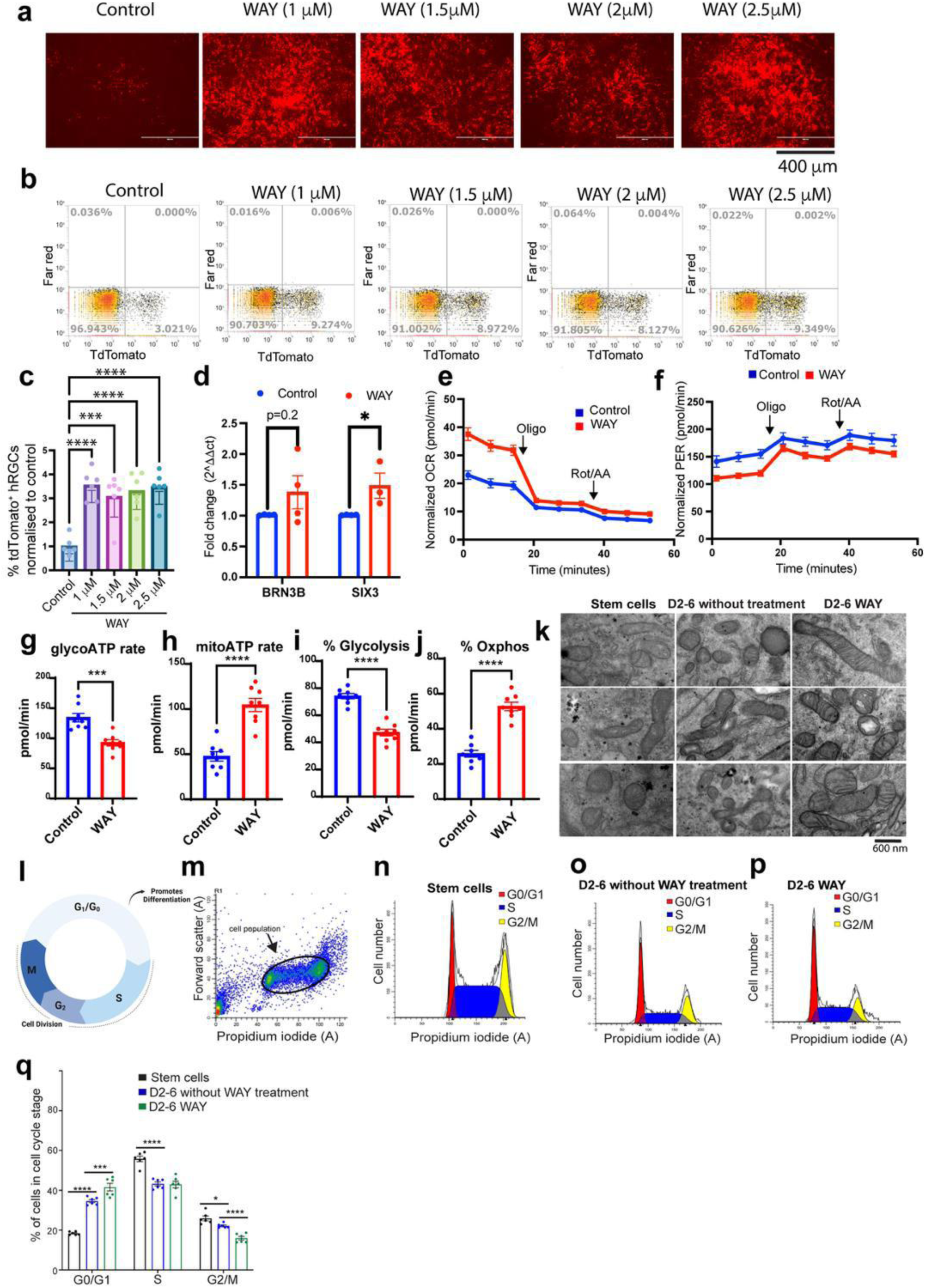
WAY-100635 reprograms metabolism to enhance hRGC differentiation. (**a**) Live-cell fluorescence of the BRN3B-TdTomato positive hRGCs on day 32 of differentiation, treated with the indicated concentrations of WAY during day 2-6. (**b**) Representative flow-cytometry plots and (**c**) quantification of percentage of TdTomato-positive hRGCs normalized to control differentiation from above samples (one-way ANOVA with Dunnett’s post-hoc test, ***P < 0.001, ****P < 0.0001; n = 6). (**d**) qRT-PCR analysis of the RGC transcription factors BRN3B and SIX3 after day 2 - 6 WAY treatment (ΔΔCt vs. GAPDH and control; unpaired two-tailed t-test, *P < 0.05; n = 3 - 4 biological replicates, each measured in triplicate). (**e, f**) Cell number normalized oxygen-consumption rate (OCR) and proton-efflux rate (PER) from the Seahorse ATP-rate assay on day 7 of hRGC (H7-hESC) differentiation cultures with or without WAY treatment (days 2–6). (**g - j**) Derived metabolic parameters - GlycoATP rate, MitoATP rate, percentage of glycolysis and OXPHOS, demonstrating a glycolysis-to-OXPHOS shift after WAY treatment (Student’s t-test, ***P < 0.001, ****P < 0.0001; n = 8). (**k**) Transmission-electron micrographs revealing elongated, cristae-rich mitochondria in WAY-treated cells (23 000×; scale bar, 600 nm). (**l**) Schematic of cell-cycle progression and G1/G0 arrest that promote differentiation. (**m**) Representative propidium-iodide flow plot of day-7 cultures indicating G0/G1, S, and G2/M fractions. (**n - p**) Cell-cycle profiles of undifferentiated H7-hESCs (**n**), day-7 untreated hRGC differentiation cells (**O**), and day-7 WAY-treated cells (**p**). (**q**) Percentage of cells at different cell-cycle phase across conditions (unpaired two-tailed t-test, *P < 0.05, ***P < 0.001, ****P < 0.0001; n = 6). Error bars are SEM.

To pinpoint the metabolic signature that underlies the permissive day 2–6 window, we interrogated bioenergetics on differentiation day 7. A 4-day pulse of WAY-100635 elevated basal oxygen-consumption rate in the seahorse ATP rate assay, which reverted to control values after electron-transport-chain blockade, indicating enhanced mitochondrial respiratory capacity (**Fig. 5e**). Proton-efflux rate (PER), an index of glycolytic acidification, remained significantly lower in the Day 2-6 WAY treated cultures (**Fig. 5f**). Deconvolution of total ATP production confirmed a sharp decline in glycoATP and %Glycolysis (**Fig. 5g, i**) with a reciprocal increase in mitoATP and %OXPHOS (**Fig. 5h, j**). Ultrastructurally, WAY-exposed differentiation culture cells on day 7 harbored elongated mitochondria with densely packed cristae, morphological hallmarks of high-efficiency oxidative phosphorylation (**Fig. 5k**). Collectively, these data show that a brief pharmacologic cue enforces an early glycolysis-to-OXPHOS switch, the metabolic milestone of neuron specification^48,49^, thereby mechanistically linking mitochondrial maturation to the amplified RGC yield.

Enhanced RGC differentiation indicates WAY treatment induces G₀/G₁ arrest in the progenitor stage as G₁ residency biases cells toward differentiation^50^ as demonstrated in **Fig. 5l**. To test this, we quantified cell-cycle distribution in the RGC differentiation culture post day 2-6 WAY treatment by propidium-iodide flow cytometry method^51^. Undifferentiated hESCs were S-phase-enriched (**Fig. 5m, n, q**). By day 7, differentiating cultures accumulated in G₀/G₁ with concomitant reduction in the S and G2/M phase, an effect magnified by early WAY exposure (**Fig. 5o, p, q**), consistent with metabolic checkpoint mediated exit from proliferation. Thus, our data show a brief surge in mitochondrial biogenesis early in development triggers mitochondrial maturation, persistent OXPHOS dominance, enforces G₁ arrest, and accelerates RGC lineage specification. This result agrees with the developmental aspect of neuronal maturation as is promoted by mitochondrial maturation and OXPHOS metabolism^52^. Thus, WAY mediated metabolic-cell-cycle coupling offers a unifying framework whereby transient pharmacologic cues can amplify stem-cell derived RGC yield, expediting a developmental outcome and pre-installing reversible stress-resilient programs that may confer in vivo neuroprotection.

### WAY protects RGCs and promotes axon regeneration to reestablish eye-to-brain connectivity with no systemic toxicity

To test the in vivo efficacy of WAY, we employed two models of optic neuropathy: an acute optic nerve crush (ONC) model and a chronic high intraocular pressure (IOP) glaucoma model induced by anterior chamber microbead injection. We evaluated the in vivo efficacy of WAY using two daily intraperitoneal (i.p.) dosing regimens: a preconditioning paradigm, in which animals were treated starting 5 days prior to optic nerve crush (ONC; Day 0) and continued until sacrifice, and a post-treatment regimen designed to mimic a clinical scenario, with dosing initiated 4 hours after ONC and continued daily thereafter (**Fig. 6a**). The 4-hour post-injury timepoint was selected based on prior evidence that irreversible neurodegenerative signaling cascades are triggered within 6 hours of ONC^53^. We found that the RGC count in naïve retina and ONC paradigm consistently induces 60 - 80% RGC loss within one week (**Fig. 6b, c**), aligning with previous reports^54,55^ providing a robust quantitative platform for evaluating neuroprotection. Daily i.p. administration of WAY, initiated prior to ONC until sacrifice, significantly preserved RGC survival at 6 days post crush (dpc) and to a lesser extent at 14 and 35 dpc (**Fig. 6a -c**). Interestingly we also have seen moderate but progressive RGC loss in the contralateral eye as obsrved by others for unilateral ONC^56,57^, and significant protection by systemic WAY treatment (**Fig. S4a, b**). Post-injury treatment alone was also effective, preserving RGCs 17 days after unilateral left ONC (**Fig. 6d - e**), thereby modeling a clinically relevant therapeutic window. Notably, despite the expected decline in efficacy at later timepoints in this severe model, the level of protection observed is likely sufficient for chronic glaucoma, where axonopathy progresses more slowly.

**Fig. 6.**
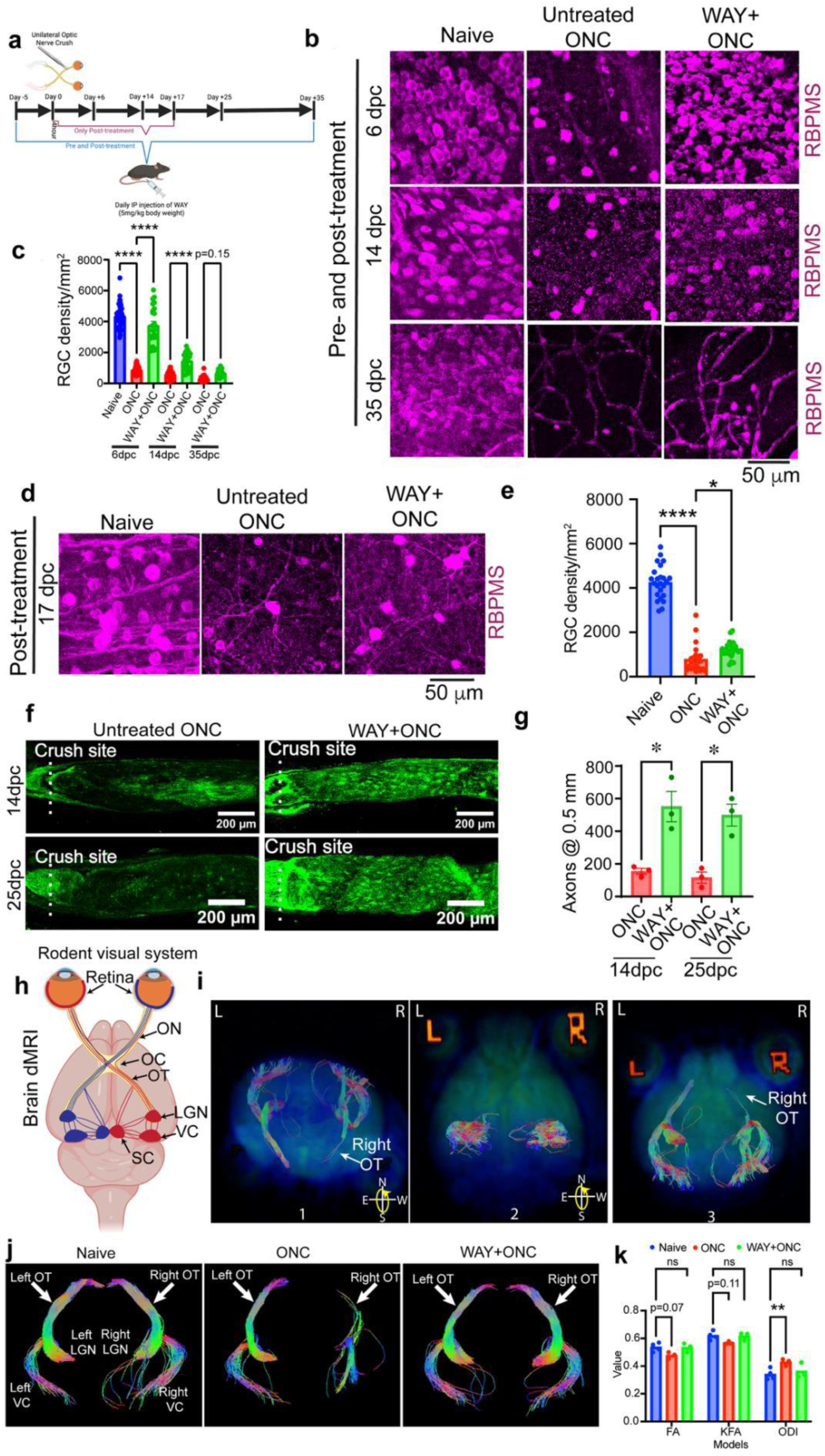
WAY-100635 preserves RGCs and restores eye-to-brain connectivity after optic-nerve crush. (**a**) Experimental design. C57BL/6J mice received unilateral left optic-nerve crush (ONC, day 0) and daily intraperitoneal WAY (5 mg/kg) either as 5 d before optic nerve crush (pre-treatment) and continuing for 6, 14, or 35 day post-crush (dpc) or starting 4 h post-injury (post-treatment) and continuing until the indicated dpc. (**b**) Representative confocal micrographs (40×/1.3 NA) of RBPMS-labelled RGCs in retina from pre-treated animals at 6, 14, and 35 dpc. (**c**) Quantification of RGC density (cells/mm^2^) from peripheral and central ONC retina for the conditions in **b** (two-way ANOVA with Tukey, ****P < 0.0001; n = 24 - 48 images from 3 - 6 mice/group). (**d, e**) Images and quantification of RGCs from ONC retina (peripheral and central) after 17 dpc when animals received daily WAY starting 4 h post-injury (two-way ANOVA with Tukey, ****P < 0.0001, *P < 0.05; n = 24 images from 3 mice/group). (**f**) GAP-43 immunostaining of longitudinal optic-nerve sections from crushed nerve at 14 and 25 dpc under pre-treatment condition; stitched mosaics show regenerating/protected axons 0.5 mm distal to the crush. (**g**) Axon counts for **f** (two-way ANOVA with Tukey, *P < 0.05; n = 3 mice/group). (**h**) Diagram of murine visual projections (∼90 % contralateral). (**i, j**) Diffusion-MRI tractography superimposed with the B0 image extracted from the diffusion-weighted imaging (DWI) dataset (**i**) and tractography only (**j**) of the optic tract (OT), lateral geniculate nucleus (LGN), and visual cortex (VC) in naïve (no treatment or injury), ONC, and WAY-treated ONC mice (pre-treatment paradigm; scan at 11 - 12 dpc). (**k**) Microstructural metrics - fractional anisotropy (FA, DTI model), kurtosis FA (KFA, DKI), and orientation-dispersion index (ODI, NODDI) for right OT region. Significant degeneration in ONC versus naïve is absent in WAY-treated mice (two-way ANOVA with Tukey, **P < 0.01; n = 3 - 4 mice/group). Error bars are SEM.

To determine whether the preserved RGCs retain the capacity for axonal regeneration, we quantified regenerating axons using GAP-43 immunohistochemistry on transverse optic nerve sections, as it marks the growing axons^58,59^. High-resolution confocal imaging with validated GAP-43 antibodies^60^ revealed minimal regeneration in untreated ONC nerves, whereas WAY-treated animals showed robust axon regrowth at 14 and 25 dpc (**Fig. 6f, g**). Importantly, the pre-crush zone displayed stronger GAP-43 signal than the post-crush segment, suggesting that the subset of RGCs preserved by WAY retains regenerative competence or that their axons were protected from degeneration due to continued treatment. This axon staining pattern and regenerating axonal count range agree to that reported in protective paradigms involving lens injury, or axon growth promoting gene perturbations^60–62^.

Crucially, axonal regeneration closely agrees with the extent of RGC soma preservation. Although RGC counts declined between 14 and 35 dpc, they remained significantly higher in WAY-treated animals than in controls (**Fig. 6b - c**). Consistently, axon counts at 14 and 25 dpc were comparable and markedly elevated relative to untreated ONC animals (**Fig. 6f, g**). These findings suggest that WAY not only prevents RGC degeneration but also supports long-range axonal growth required to establish lost connectivity to the brain. To support direct target engagement of WAY with 5-HT1A in RGCs, we first confirmed receptor expression in human and mouse RGCs and assessed compound accessibility across the blood-retinal barrier. Immunofluorescence revealed strong plasma membrane localization of 5-HT1A in human stem cell-derived hRGCs (**Fig. S5a**), corroborated by western blot of whole-cell lysates (**Fig. S5b**). In mouse retina, confocal immunohistochemistry imaging showed robust 5-HT1A expression in the ganglion cell layer (GCL), which remained stable in surviving RGCs 25 days after ONC (**Fig. S5c**). To quantify receptor levels, we purified RGCs from mouse retina using magnetic-activated cell sorting (MACS), the method reported to have minimal perturbation of native transcriptomic and proteomic profiles^12,63^ and obtained nearly 90% purity (**Fig. S5d, e**). Isolated RGCs exhibited sustained 5-HT1A expression 24 h post-ONC (**Fig. S5f**), supporting continuous target availability in disease context. WAY crosses blood-brain barrier following intraperitoneal injection in mice^64^. To assess bioavailability in retina, we performed mass spectrometry following a single intraperitoneal injection of WAY. Chromatographic peaks confirmed compound presence in both plasma and retina, with retinal levels peaking at 4 h and declining by 24 h (**Fig. S6a, b**), consistent with daily dosing rationale. Given the chronic nature of glaucoma, systemic safety is essential. Mice treated daily with WAY (5 mg/kg, i.p.) for 30 days exhibited no structural abnormalities in kidney or liver by hematoxylin and eosin (H&E) staining. In particular, we observed no glomerular lesions (**Fig. S6c,** yellow arrows) nor necroinflammatory changes surrounding the hepatic central vein (**Fig. S6d**, yellow arrows) that marks systemic toxicity^64,65^.Mice also maintained normal behavior, body weight, and activity throughout the treatment period. Together, these data confirm 5-HT1A expression in RGCs, efficient retinal penetration of WAY, and a favorable safety profile, supporting on-target efficacy in both in vitro and in vivo models.

Neurodegeneration of brain visual centers is a common and debilitating consequence of glaucoma progression^66,67^, yet no existing therapy has demonstrated preservation or regeneration of RGC axons into brain targets, an essential prerequisite for functional neuroprotection. To build upon our evidence of optic nerve axon regeneration (**Fig. 6f - g**), we next asked whether WAY treatment supports long-range axonal preservation or regeneration into retino-recipient brain regions. We employed double-blinded diffusion MRI (dMRI) analysis to assess white matter integrity in vivo. All imaging and computational analyses were performed under identity-masked conditions. Given that ∼90% of mouse RGCs project contralaterally^68^ as demonstrated in **Fig. 6h**, we performed unilateral left ONC and focused on the right-side visual structures. In untreated animals, B0 (baseline image with no diffusion weighting) and tractography overlays confirmed optic tract (OT) degeneration predominantly in contralateral to the crushed eye, consistent with expected pathophysiology (**Fig. 6i; Supplementary Movie 1**). Using fully automated, unsupervised tractography algorithms^69–71^, we found that ONC resulted in dramatic loss of fiber connectivity in the right OT and downstream visual centers by 11–12 days post crush. In contrast, animals treated with WAY 5 days prior to ONC until imaging exhibited robust preservation of fiber tracts along the same pathway (**Fig. 6j**), aligning with our earlier immunohistochemical evidence of axon regeneration. To quantitatively evaluate microstructural integrity, we analyzed diffusion-weighted image volumes using multiple tensor models, extracting key structural metrics including Fractional Anisotropy (FA)^72^, Kurtosis Fractional Anisotropy (KFA)^73^, and Orientation Dispersion Index (ODI)^74^ in the right OT. FA and KFA, both indicators of axonal integrity and microstructural complexity^75^ were reduced in ONC animals but preserved in the WAY-treated cohort (**Fig. 6k**). Conversely, ODI, which increases with disorganized fiber orientation, was elevated in ONC mice but remained low in treated animals, further indicating structural preservation (**Fig. 6k**).

Taken together, these blinded, quantitative dMRI results independently confirm that WAY not only protects RGC somas and proximal axons but also maintains retino-brain connectivity across the visual axis. This represents, to our knowledge, the first demonstration of pharmacological axon preservation into brain visual centers in a glaucoma-relevant model, a critical step toward realizing functional neuroprotection in human disease.

### In vivo WAY restores RGC activity post injury and provides glaucoma neuroprotection

Building on our findings that WAY promotes axon regeneration into central visual targets, we next tested whether this structural rescue translates into functional preservation of RGC activity under neurodegenerative conditions. Using flash electroretinography (ERG) under rod-saturating green light (Diagnosys LLC), we recorded the photopic negative response (PhNR), a cone-driven RGC-specific readout, alongside the a-wave and b-wave components reflecting upstream retinal function^76–78^, under the post treatment only condition (**Fig. 7a, b**). In ONC-injured mice, the affected eye exhibited a marked reduction in amplitude and delay in the PhNR, consistent with RGC dysfunction. Post-injury treatment with WAY significantly restored PhNR amplitude and timing (**Fig. 7b, c**), indicating functional rescue of RGCs even after axonal damage.

**Fig. 7.**
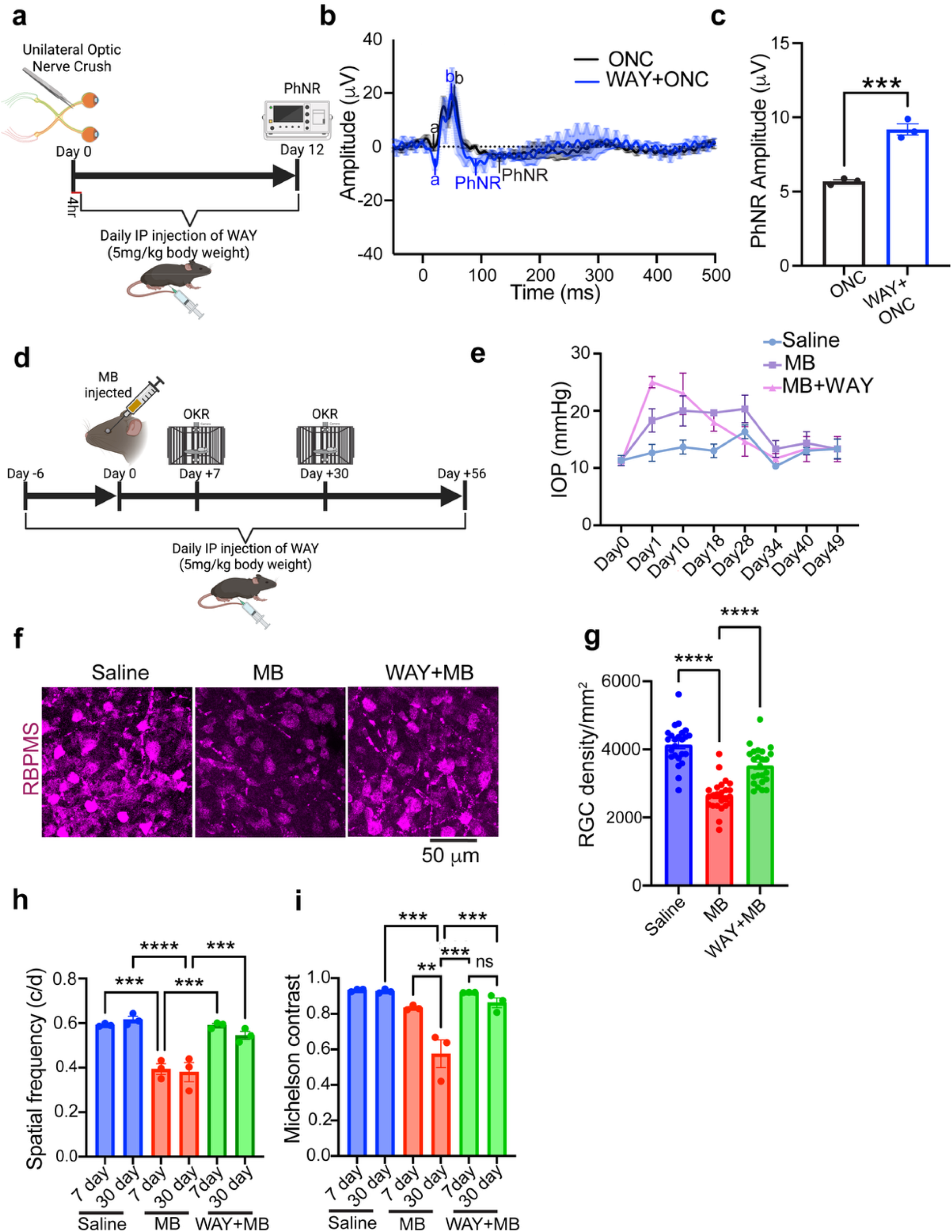
**WAY-100635 restores RGC function after optic nerve injury and protects against chronic IOP elevation**. (**a**) Experimental timeline for unilateral ONC in C57BL/6J mice followed by daily intraperitoneal WAY (5 mg/kg) beginning 4 h post-injury; photopic negative response (PhNR) recordings were obtained 12 days post-ONC. (**b**) Representative flash-ERG traces (white flashes on a rod-saturating green background) illustrating PhNR peaks; each trace is the average of nine recordings (three technical replicates per eye, three eyes). (**c**) PhNR amplitude under the indicated conditions (n = 3 mice/group). Student’s t-test, ***p < 0.001. (**d**) Schematic for chronic glaucoma model: unilateral microbead (MB) occlusion on day 0 (2 µL of 1 µm + 2 µL of 6 µm polystyrene beads) or PBS (saline) into the anterior chamber, followed by daily WAY dosing (5 mg/kg). Visual acuity (VA) and contrast sensitivity (CS) were assessed with OptoMotry on days 7 and 30 (see panels H-I). (**e**) Intraocular pressure (IOP) profile measured by rebound tonometry: sustained elevation for ∼4 weeks with subsequent return toward baseline. (**f**) Representative confocal images of RBPMS-immunolabeled RGCs 56 days after MB induction ± WAY. (**g**) RGC density (cells/mm^2^) quantified from 24 retinal fields (peripheral and central) from MB occluded eye (n = 3 mice/group). Two-way ANOVA with Tukey post-hoc, ****p < 0.0001. (**h-i**) Functional vision outcomes. (**h**) VA (cycles/deg) determined by head-tracking to drifting gratings of increasing spatial frequency. (**i**) CS (Michelson contrast) determined by head-tracking to gratings of decreasing contrast. Measurements were performed under blinded conditions for both clockwise and counter-clockwise rotation; two-way ANOVA with Tukey correction, **p < 0.01, ***p < 0.001, ****p < 0.0001; n = 3 mice/group. Data are mean ± SEM.

Given these robust effects in the ONC model, we next asked whether WAY preserves RGC structure and function under chronically elevated IOP, a more clinically relevant model of glaucoma. We induced unilateral ocular hypertension using microbead (MB) injection into the anterior chamber, a well-established model that elevates IOP without ischemic injury or off-target retinal damage^79,80^. We saw IOP elevation to ∼20 mmHg on MB injected eyes and persisted for approximately 4 weeks before returning to baseline (∼12 mmHg) (**Fig. 7d, e**) consistent with prior reports^81^. Interestingly, WAY-treated mice showed a slightly accelerated IOP elevation and resolution (**Fig. 7e**). Previous studies have shown that despite IOP returning to baseline within a month following a single unilateral MB injection, RGC degeneration continues for at least 8 weeks^82^. Consistent with these findings, we observed that even transient IOP elevation was sufficient to cause significant RGC loss by 8 weeks in the untreated animals. Remarkably, daily treatment with WAY (6 days before MB to day of measurements, i.p.) robustly preserved RGC density (**Fig. 7f, g**), reinforcing its neuroprotective efficacy in both acute (ONC) and chronic (IOP-induced) models of optic neuropathy. To evaluate visual function, we used the OptoMotry system to measure visual acuity and contrast sensitivity. Visual acuity declined within 7 days post-MB injection and plateaued by 4 weeks, a trajectory recapitulating adaptive changes driven by the contralateral eye^9^. WAY treatment significantly mitigated acuity loss (**Fig. 7h**), indicating functional benefit beyond anatomical protection. On the other hand, contrast sensitivity exhibited a progressive decline between days 7 and 30 in IOP induced untreated mice, which was substantially attenuated in the WAY group (**Fig. 7i**).

Together with the axonal regeneration data, these results demonstrate that WAY confers robust and durable neuroprotection by preserving both the structure and function of RGCs across distinct models of optic neuropathy, supporting its translational potential as a glaucoma therapy.

## DISCUSSION

### WAY-100635 as a first-in-class multi-pathway neuroprotective agent

This study establishes the 5-HT1A receptor antagonist WAY as a promising therapeutic candidate for glaucoma. Integrating mechanistic insights from human pluripotent stem cell-derived RGCs (hRGCs) with in vivo anatomical and functional validation, we demonstrate that WAY initiates a multifaceted, reversible neuroprotective program, beginning with cAMP signaling, mitochondrial biogenesis and metabolic reprogramming, culminating in long-distance axonal regeneration into central visual targets and preservation of vision. Restoration of eye-to-brain connectivity following optic nerve injury has long represented a frontier challenge in glaucoma and other optic neuropathies. Foundational studies using peripheral nerve grafts showed that appropriate extrinsic cues can drive RGC axon regeneration through the injured optic nerve and into the superior colliculus^83,84^, establishing the principle that intrinsic growth programs can be reactivated with the right stimuli. Since then, genetic manipulations such as Pten or Socs3 deletion^85^, Sirt1 overexpression^86^, and activation of transcription factors like Sox11^87^ or trophic factors like CNTF^88,89^ have demonstrated regenerative capacity in experimental systems. However, these approaches remain invasive, difficult to control temporally, and largely limited to research settings. In contrast, our study introduces a systemically deliverable small molecule with a favorable human safety profile that engages reversibly multiple neuroprotective and regenerative pathways. WAY enables robust axon protection and regrowth, thereby reestablishing eye-to-brain connectivity in vivo. These findings position WAY as a compelling pharmacological candidate for advancing neuroprotective therapy in glaucoma and potentially other central nervous system injuries.

### Metabolic reprogramming by WAY supports both RGC differentiation and regeneration

Our data reveal a dual and stage-specific benefit of WAY-mediated metabolic reprogramming. Early treatment during the progenitor phase enhances mitochondrial maturation, promotes oxidative phosphorylation (OXPHOS), accelerates cell-cycle exit, and increases the yield of differentiated RGCs (**Fig. 5**). In contrast, treating mature WT or OPTN^E50K^ hRGCs restores mitochondrial health but favors a shift toward aerobic glycolysis (**Fig. 4**). These distinct metabolic outcomes carry two important implications.

Metabolic reprograming with enhanced mitochondria dependent metabolism are important for stem cell differentiations including cardiomyocytes, and neurons^52,90^. However, we still lack a safe pharmacological compound that will achieve this goal to overcome the translational barrier. First, pharmacologically enhancing RGC differentiation with a safe compound like WAY could transform stem cell-based cell replacement strategies for late-stage glaucoma. A key limitation in such therapies is generating sufficient numbers of lineage-committed RGCs while minimizing undifferentiated or proliferative cells that carry teratoma risk when transplanted^91,92^. Our finding that WAY-treated progenitors exit the cell cycle earlier and yield more RGCs suggests that fewer, metabolically primed progenitors may be sufficient for transplantation, reducing tumorigenic risk while improving therapeutic efficiency. Second, in mature RGCs, the induction of healthy mitochondria alongside an aerobic glycolytic shift appears tailored for axonal regeneration. Notably, healthy mitochondria can support aerobic glycolysis without promoting excessive OXPHOS, thereby limiting oxidative stress^93,94^. Aerobic glycolytic state likely supports the synthesis of lipids and proteins essential for membrane expansion and cytoskeletal remodeling during axon outgrowth^95^. Indeed, our diffusion MRI findings show that WAY-treated ONC mice maintain near-normal axonal complexity in the optic tract 11–12 days post-injury, despite significant RGC loss by day 14 (**Fig. 6b, c, f-k**). This preserved complexity suggests that surviving RGCs not only regenerate axons but also initiate axon sprouting and branching, a process reminiscent of developmental arborization and synaptogenesis in brain visual centers^96,97^.

Together, these results position WAY as a context-dependent modulator of RGC metabolism: promoting OXPHOS and differentiation in progenitors while enabling glycolysis-driven anabolic growth and axonal remodeling in mature neurons. This dual mechanism supports both the generation and long-distance regeneration of RGCs, offering a powerful strategy for vision restoration in glaucoma.

### Context-dependent effects of 5-HT1A modulation on RGC neuroprotection

While our findings support the 5-HT1A receptor antagonist WAY as a promising neuroprotective agent for glaucoma, prior studies have reported protective effects of 5-HT1A agonists such as 8-OH-DPAT (DPAT), leading to apparent discrepancies in the field^98,99^. These differences likely stem from context-dependent effects of 5-HT1A signaling on RGC physiology and survival under varying experimental conditions.

Mechanistically, 5-HT1A is a Gi/o-coupled GPCR whose activation suppresses adenylyl cyclase and reduces intracellular cAMP. In contrast, antagonism by WAY relieves this inhibition, leading to cAMP elevation, a second messenger known to support neuronal survival and axonal regeneration^29,30,100^. Our data demonstrate that WAY treatment reversibly increases cAMP, enhances mitochondrial biogenesis, and reduces apoptosis in OPTN^E50K^ and WT hRGCs. Conversely, DPAT treatment decreased cAMP (**Fig. 1**) and increased apoptosis in the OPTN^E50K^ hRGCs (**Fig. 3j**), suggesting an opposing, contextually maladaptive effect under disease-relevant conditions. Interestingly, selective seretonin reuptake inhibitors (SSRIs), which increase serotonin concentration in synaptic cleft, are linked to higher glaucoma risk^101^, likely due to overstimulation of 5-HT1A receptors. Several factors may account for the divergence in reported effects. First, it has been shown that enhanced RGC excitability within 2 weeks of elevated IOP is protective, which declines after 4 weeks with axonal degeneration^9,102,103^. However, the above studies show that while DPAT suppresses RGC excitation at 4 weeks of IOP elevation, WAY rescues it^98,99^, which supports WAY as protective and DPAT as degenerative. This suggests DPAT’s protective effects may be indirect, potentially via lowering IOP as was observed in rabbits^104^. However, WAY in our hand, did not significantly affect IOP in mouse MB induced glaucoma model, indicating its direct neuroprotective effect. Another anomaly may arise from the retrograde labeling method used in detecting RGC protection. The authors used fluorogold labeling to assess RGC survival ^99^, but this method detects retrograde transport defects (brain to eye) rather than true cell survival^5^.

Taken together, while 5-HT1A agonists may offer transient benefits under specific conditions, our mechanistic and functional data favor 5-HT1A antagonism via WAY as a more consistent and directly protective strategy for promoting RGC survival and axon regeneration in glaucoma.

### Limitations and Future Directions

Our findings reveal that WAY’s neuroprotective effect extends beyond metabolic reprogramming and likely involves a convergence of pro-survival and regenerative signaling cascades downstream of 5-HT1A antagonism. Future studies need to resolve the RGC protection versus regeneration effect of WAY and involved downstream signaling pathways. The downstream signaling cascade may involve cAMP regulation of CREB, CaMKII, and SIRT1 factors that have known RGC protective effect^86,105,106^. These pathways may collectively contribute to the robust protection and regeneration observed in both human stem cell-derived RGCs and in vivo models but their involvement yet to be established. Additionally, although our analyses focused on RGC-intrinsic effects, the contributions of non-neuronal cell types including astrocytes, microglia, and vascular cells are not yet defined. These populations could modulate inflammation, provide metabolic support, or secrete trophic factors that enhance WAY’s efficacy in vivo. Ultimately, a deeper understanding of the molecular and cellular landscape engaged by WAY will guide rational combination therapies and inform translational strategies for optic nerve regeneration and neurodegeneration more broadly.

## Conclusion

This study identifies WAY-100635 as a pharmacological agent that restores mitochondrial homeostasis, promotes RGC differentiation, and preserves visual system integrity across multiple models of glaucoma. By antagonizing 5-HT1A receptor, WAY engages a cascade of neuroprotective mechanisms that unify metabolic support, transcriptional plasticity (PGC-1α), and axonal regeneration. These findings open new avenues for repurposing serotonergic modulators in neurodegenerative diseases and highlight the power of targeting intrinsic neuronal pathways for durable vision preservation.

## Resource Availability

The resources used in this study have beeen detailed in the method and Supplemental table 1. Requests for further information and resources should be directed to and will be fulfilled by the lead contact, Arupratan Das.

## Materials Availability

This study did not generate new unique reagents.

**Table 1.**
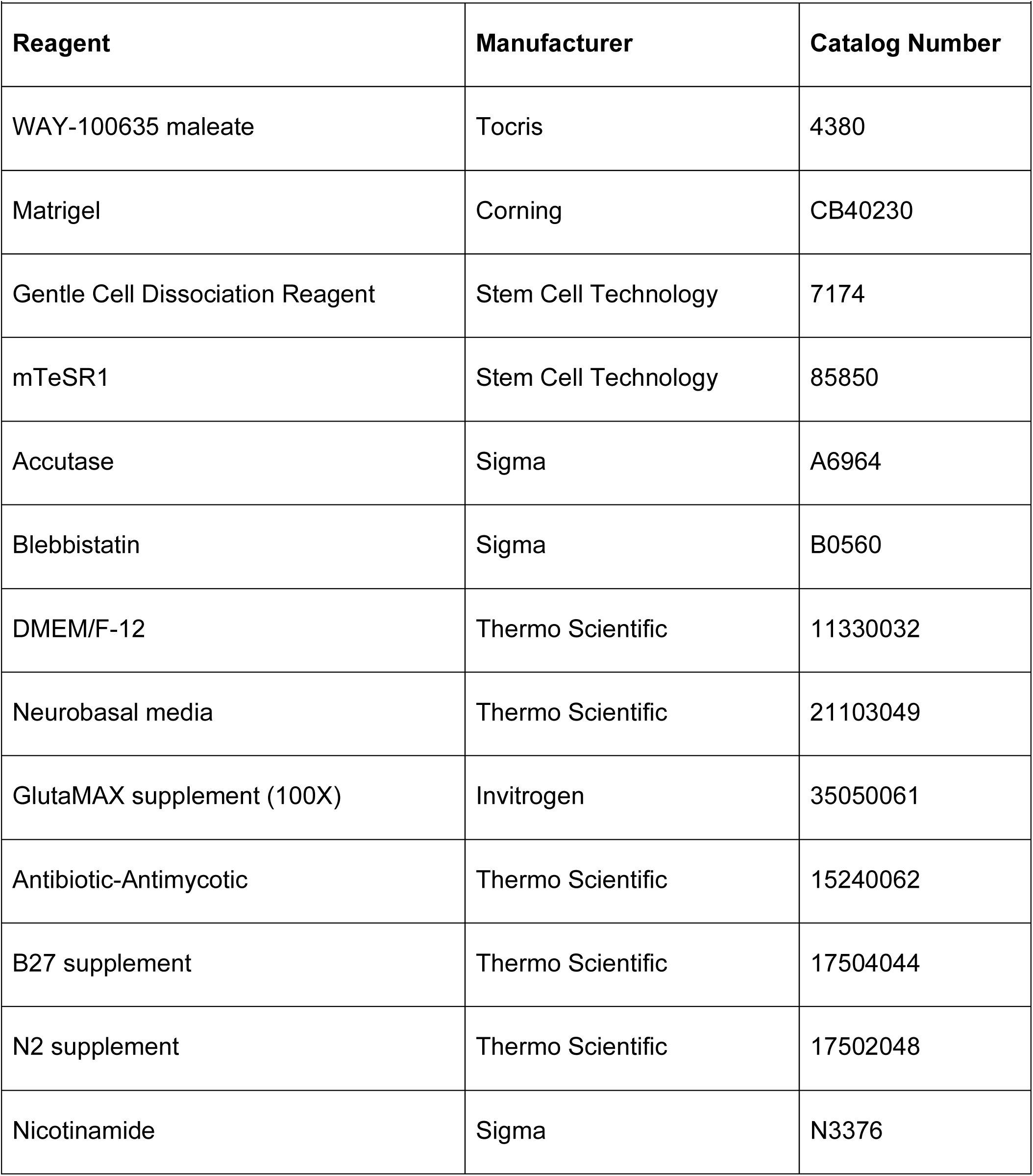
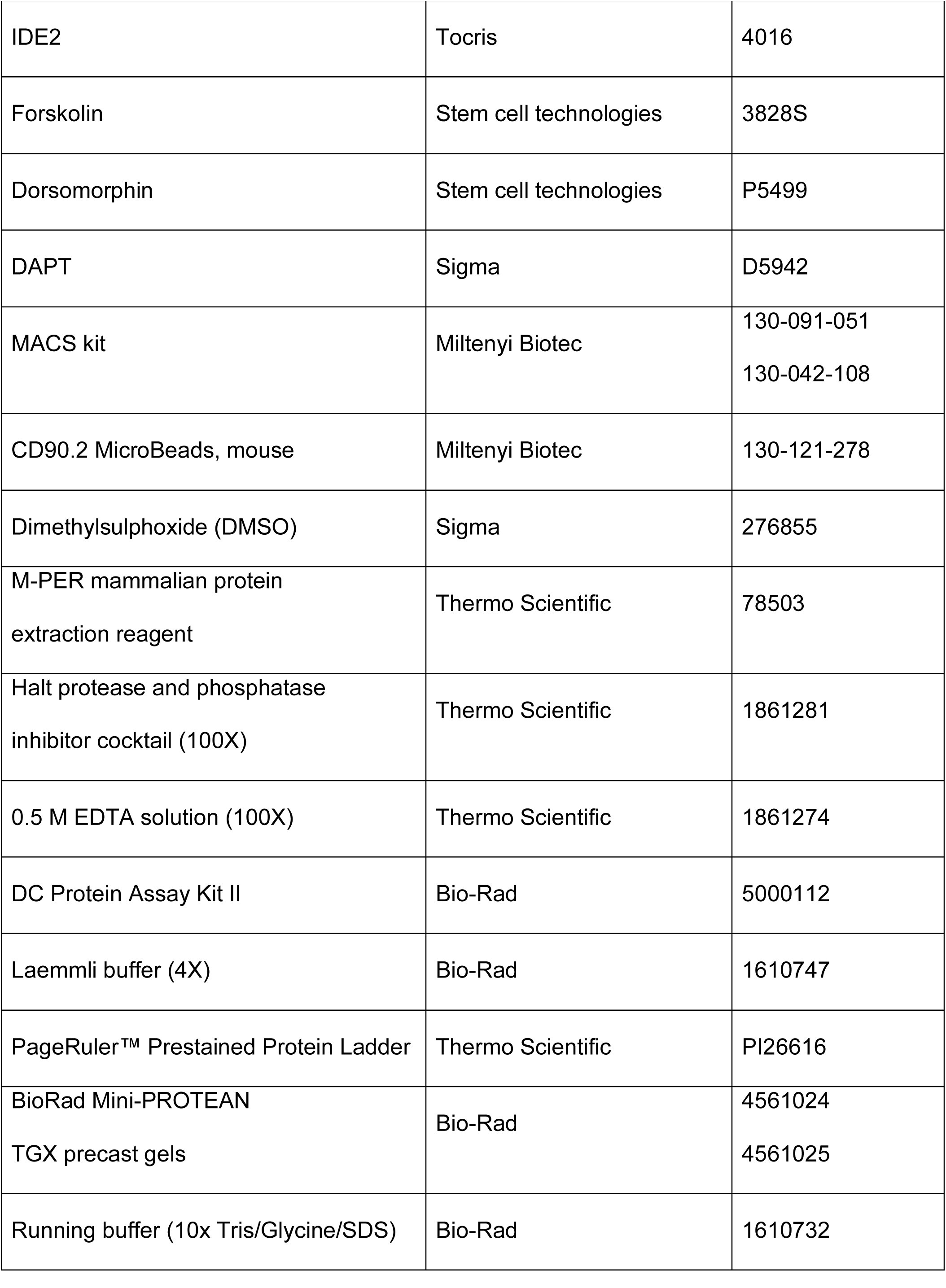
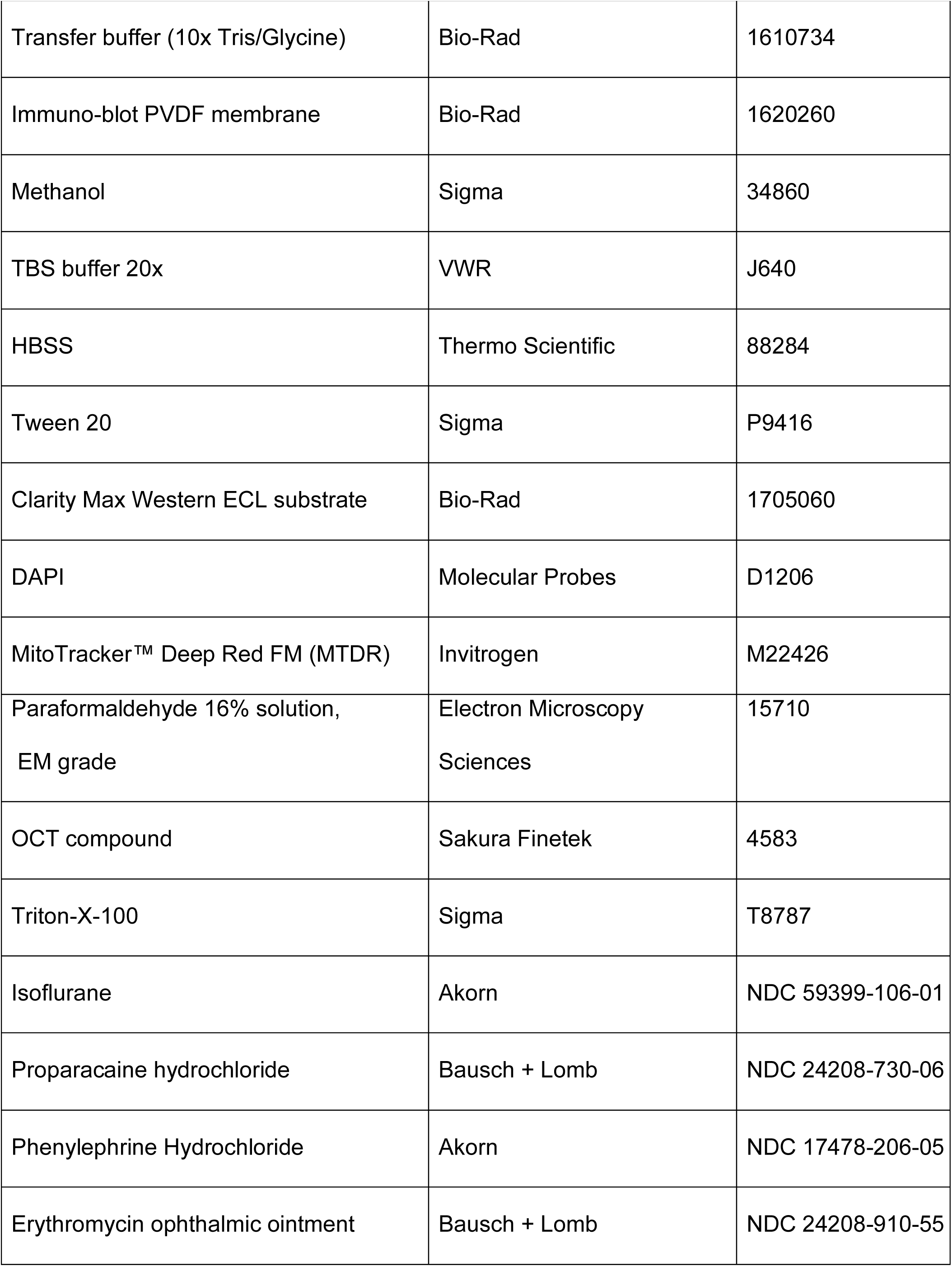
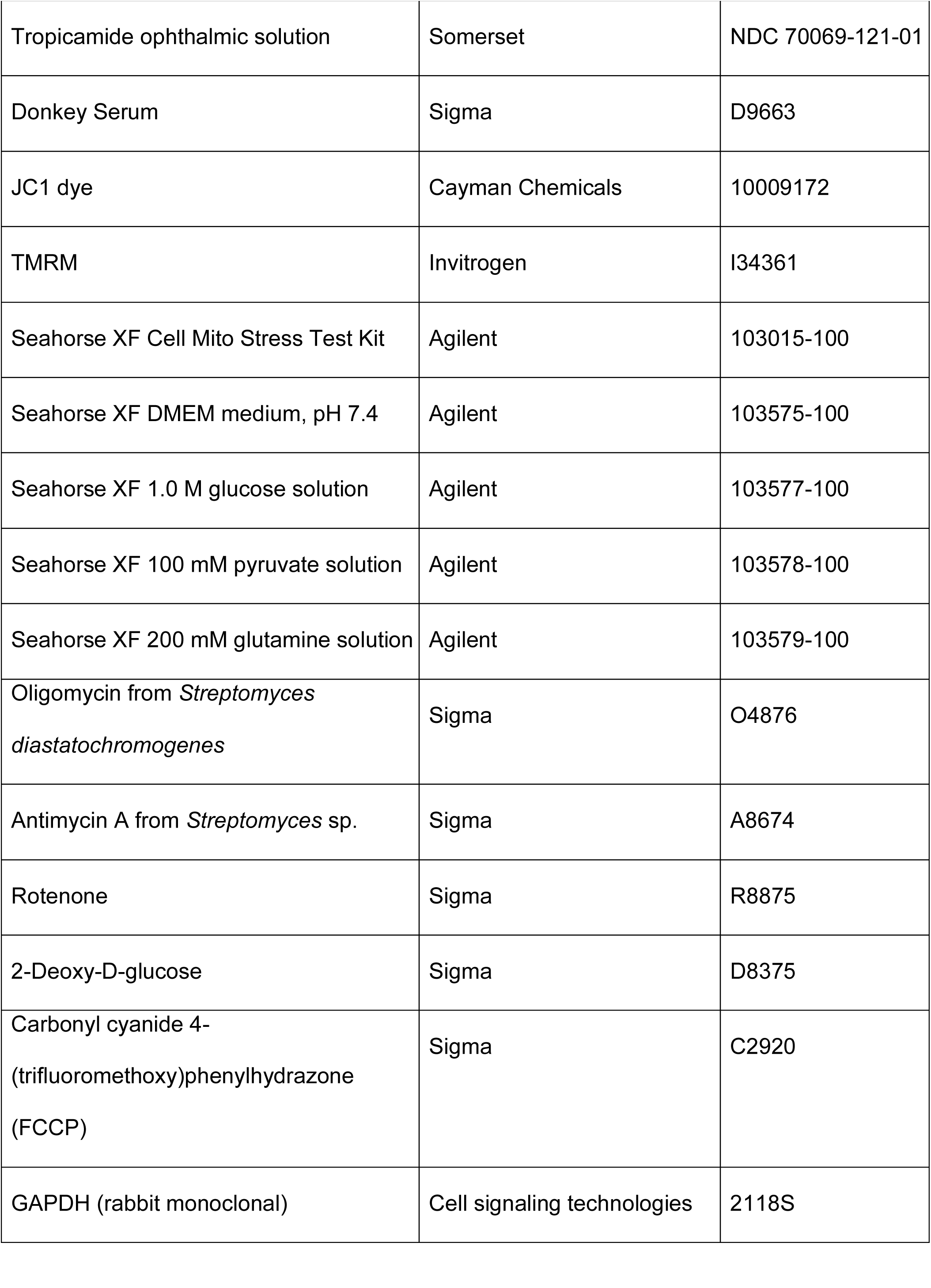
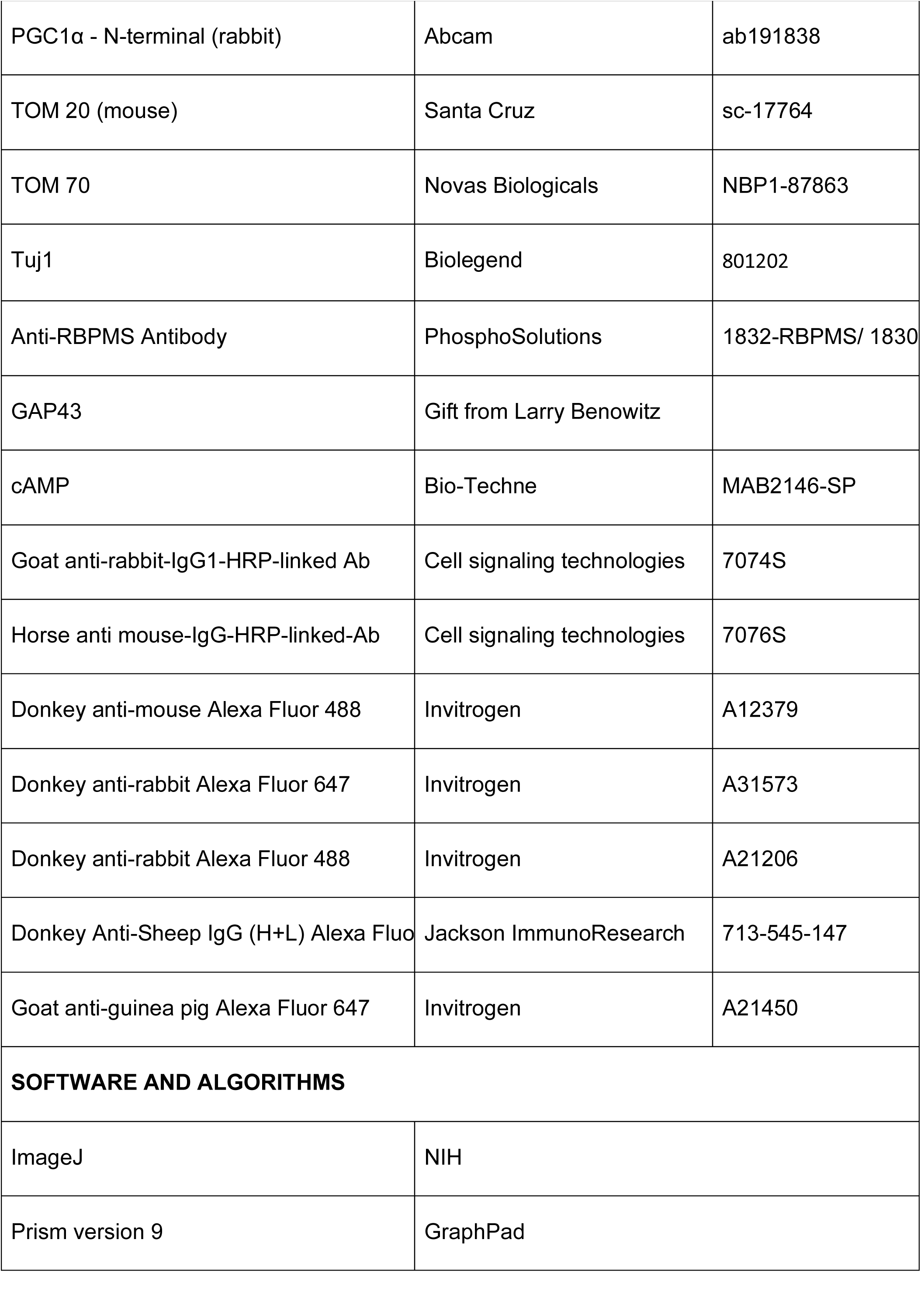
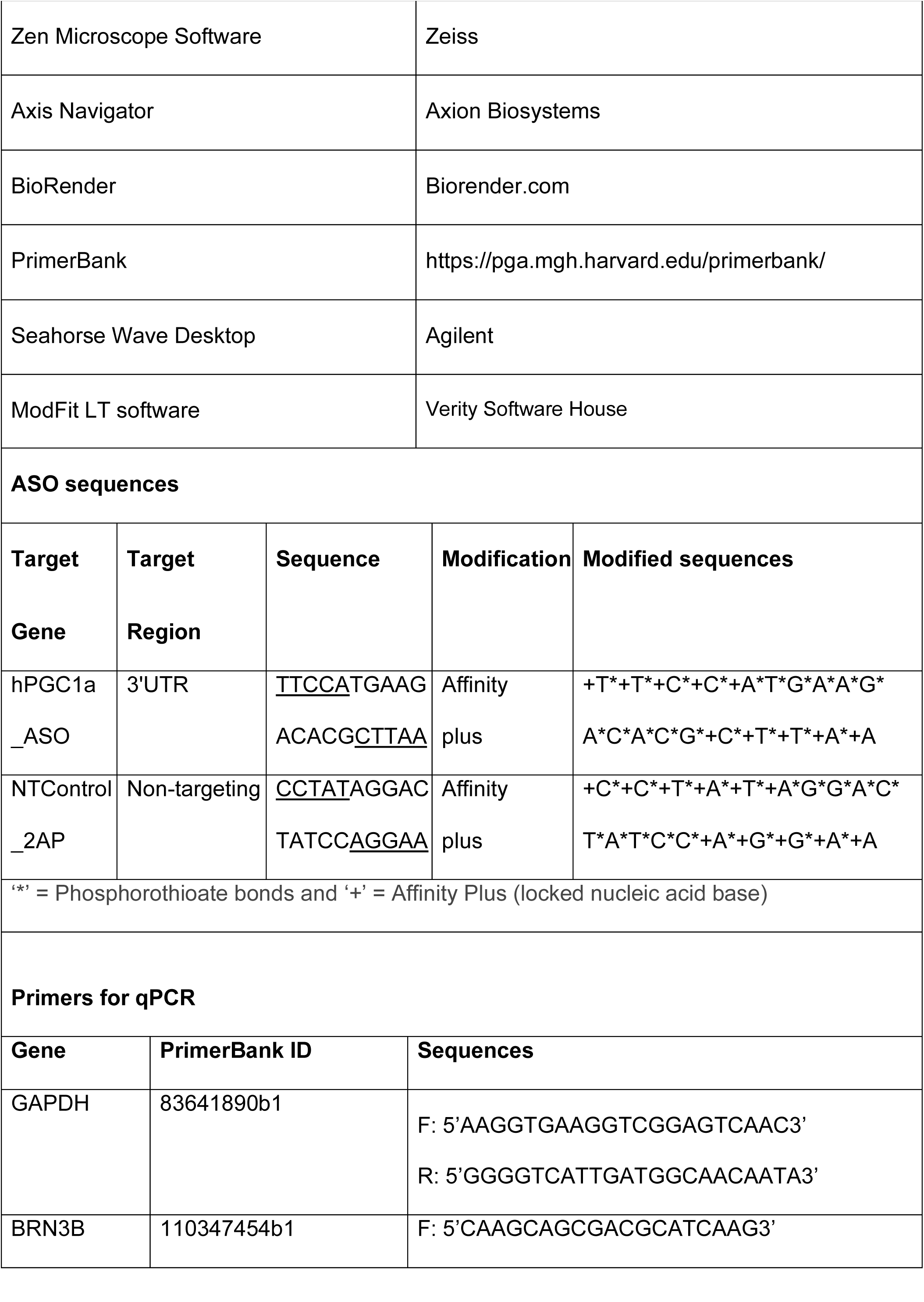
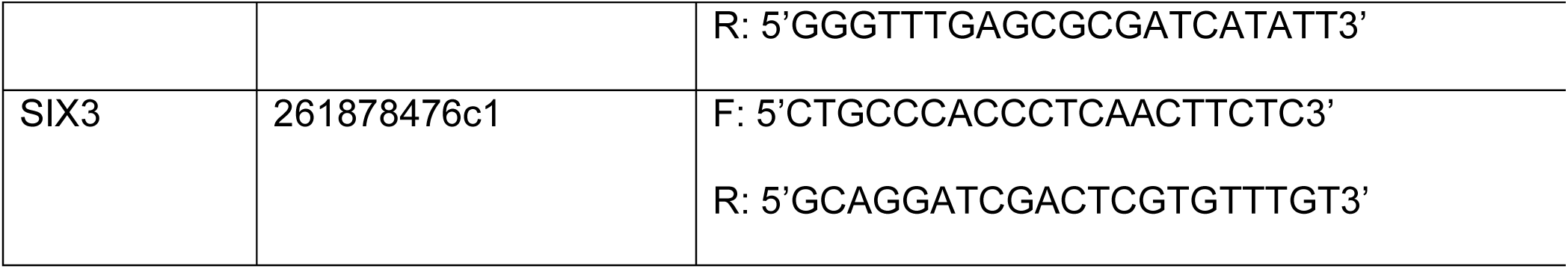
List of reagents, software, antibodies.

## Data and code availability

The data used to generate plots will be deposited in the publicly available repository and available through contacting the lead contact. All uncropped images for western blots are presented in the Supplementary Fig. 7 and flow cytometry gating steps are presented in the Supplementary Fig. 8.

This paper does not report original code

## Supporting information

Superimposed B0 image and tractography orientations depicting right OT degeneration post unilateral left ONC, related to Fig. 6

## Acknowledgements

We thank the Hypoxia Core Facility at the Indiana University Cooperative Center for Excellence in Hematology (U54 DK106846) for Seahorse support, and Dr. Donald Zack for insightful feedback. Transmission-electron microscopy was performed with Derrick Gray, Dr. Quyen Q. Hoang, and Yangshin Park at the Indiana University School of Medicine (IUSM) Electron Microscopy Core; MRI sample preparation and imaging was assisted by Dr. Yu-Chien Wu and Erin E. Jarvis at the IU in vivo imaging core. We are thankful to Andi R. Masters and Christine M. Bach at the Clinical Pharmacology Analytical Core (CPAC, IUSM) for mass-spectrometry analyses, and to Drew M. Brown (Histology Core, IUSM) for tissue sectioning and H&E staining.

We acknowledge Drs. Neha Mahajan, Qianyi Luo, and Ashay D. Bhatwadekar for OKR and ERG recordings; Drs. Amir R. Hajrasouliha and Sunland Gong for PhNR measurements; and Drs. Padmanabhan Pattabiraman and Avinash Soundararajan for IOP monitoring and anesthesia. Guidance in cryosectioning and staining was provided by Drs. Yoshikazu Imanishi, Sanae Imanishi, and Shimpei Takita; Dr. Weiming Mao for the acces to his light-microscope; and Dr. Timothy W. Corson, Dr. Anbukkarasi Muniyandi, and Kamakshi Sishtla assisted with cell ccycle analysis, enucleation and retinal dissection procedures.

We thank Dr. Erica Cai (Indiana Biosciences Research Institute) for antisense-oligonucleotide (ASO) design, and Dr. Alyssa Coyne (Johns Hopkins University) for guidance on gymnotic ASO delivery. Drs. Thomas V. Johnson III and Erika A. Aguzzi (Johns Hopkins University) trained us in optic-nerve crush and retinal flat-mounts; Dr. Harry A. Quigley and Elizabeth Kimball (Johns Hopkins University) assisted with the microbead glaucoma model; and Dr. Larry Benowitz and Dr. Yuqin Yin (Harvard Medical School) provided the GAP-43 antibody and quantification protocols.

Funding was provided by the NIH (R00 EY028223 to A.D.; UL1 TR002529 to S.D. and A.D., R01 NS125020 to N.W.), Research to Prevent Blindness Challenge Grant to IU Ophthalmology, Indiana CTSI, The Glaucoma Foundation, Showalter Research Trust, Indiana University School of Medicine, and the BrightFocus Foundation (RReSTORe collaborative grant). K.A. was supported in part by the Paul and Carole Stark Fellowship and a Sigma Xi Grant-in-Aid of Research.

The content is solely the responsibility of the authors and does not necessarily represent the official views of the NIH.

## Author contributions

A.D. conceived the study and supervised all aspects of the project. A.D., S.D., M.S., and K.A. designed the experiments, carried out the research, and analyzed the data. J.M. designed and executed the small-molecule screen. N.W. and J.C. processed, analyzed, and interpreted the MRI data.

## Supplemental information

Document S1. Fig. S1-S8 and Table S1

Video S1. Superimposed B0 image and tractography orientations depicting right OT degeneration post unilateral left ONC, related to Fig. 6

## Methods

Reagents and software used in this study are listed in Table S1.

### ESC maintenance

Human embryonic stem cells (H7-hESCs; WiCell, Madison, WI) were used as the RGC reporter line. These cells were genetically modified via CRISPR/Cas9 to insert a multicistronic BRN3B-P2A-tdTomato-P2A-Thy1.2 construct into the endogenous BRN3B locus, enabling selective fluorescent labeling and surface tagging of retinal ganglion cells (RGCs) as an isogenic reporter system^21^. A second CRISPR-engineered reporter line (H7-hESC (OPTN^E50K^)) carrying the glaucoma-associated E50K mutation in the OPTN gene was previously described and used by us for disease modeling^20^. All hESC lines were maintained in mTeSR1 medium (STEMCELL Technologies) on Matrigel-coated plates at 37°C in a humidified incubator with 5% CO₂. Routine passaging was performed using Gentle Cell Dissociation Reagent (STEMCELL Technologies) when cultures reached 70–80% confluency. Briefly, media were aspirated, cells were incubated with Gentle Dissociation Reagent for 5 minutes, and colonies were dissociated into small clumps via gentle trituration in fresh mTeSR1 before being reseeded onto new Matrigel-coated plates.

### hRGC differentiation and immunopurification

Stem cell colonies were dissociated into single cells using Accutase (Sigma) for 10 minutes at 37°C. Enzymatic activity was quenched by adding an equal volume of mTeSR1 medium (mT) supplemented with 5 μM blebbistatin (blebb). The cells were then centrifuged at 150 × g for 6 minutes, resuspended in mT containing 5 μM blebb, and counted using a hemocytometer. A total of 100,000 cells were plated per well of a Matrigel-coated 24-well plate. After 24 hours, the medium was replaced with fresh mT lacking blebbistatin. Another 24 hours later, the culture medium was switched to induction neural specification (iNS) medium, and directed differentiation toward retinal ganglion cells (hRGCs) was initiated using a small molecule -based protocol as previously described^22^. Differentiation was monitored by expression of the tdTomato reporter under the BRN3B promoter. Between days 45 and 55, tdTomato⁺ hRGCs were enriched using anti-THY1.2 microbeads and magnetic-activated cell sorting (MACS; Miltenyi Biotec), following previously established procedures^21,22^. Purified hRGCs were resuspended in iNS medium, counted, and plated onto Matrigel-coated coverslips, plates, or MatTek dishes for downstream analyses.

To enhance hRGC differentiation, cultures were treated with WAY-100635 at various time points and doses, as indicated in Fig. 5a - c and Supplementary Fig. 3a - c. Live cell fluorescence images of TdTomato-positive cells were captured on day 32 of differentiation using a Nikon EVOS microscope equipped with a 10×/0.25 NA objective. Following imaging, cells were collected for flow cytometric analysis to quantify the percentage of TdTomato-positive cells, representing the differentiated RGCs.

### Small Molecule Screening

Cold Matrigel (30 μl) was dispensed into each well of a 384-well plate using a Thermo Scientific Multidrop 384 and incubated overnight at 37 °C in 5% CO₂ to allow for gelation. The following day, excess Matrigel was removed by inverting the plate onto paper towels, and 50 μl of a cell suspension containing 10,000 hRGCs in iNS medium was added to each well. After a 48-hour incubation, media was aspirated using an automated vacuum system, and cells were labelled with 50 μl iNS medium containing 10 nM MitoTracker Deep Red (MTDR) for 15 minutes. Drug compounds were then dispensed using a Tecan EVO100 with a PIN tool, delivering 0.25 μl from a 1 mM stock plate to achieve a final concentration of 5 μM, and the cells were incubated for an additional 24 hours. Subsequently, the media was aspirated using an automated vacuum, and cells were treated with 15 μl Accutase per well for 20 minutes at 37 °C. To each well, 50 μl of iNS medium was added, and the cell suspension was mixed thoroughly by pipetting. The resulting single-cell suspensions were transferred using a Tecan EVO150 MCA with disposable tips into a 384-well U-bottom plate. Plates were then analyzed on a Thermo Fisher Attune NxT flow cytometer equipped with an autosampler, and MTDR fluorescence intensity was quantified to assess mitochondrial content across drug-treated conditions relative to DMSO controls.

### Mitochondrial mass measurement by Flow cytometry

Purified hRGCs were seeded at a density of 30,000 cells per well in Matrigel-coated 96-well plates and cultured for 3 days in iNS medium. To assess mitochondrial mass, cells were first labelled with MitoTracker Deep Red (MTDR), a mitochondrial membrane potential sensitive live cell dye. Following MTDR labeling, hRGCs were treated with WAY-100635 at various time points. At the end of each treatment, cells were dissociated into single-cell suspensions using Accutase, and mitochondrial content was quantified using a Thermo Fisher Attune NxT flow cytometer. Fluorescence intensity from far-red channel (MTDR) was used as a direct readout for average mitochondrial mass in individual cell.

### Cell viability and caspase activity

Stem cell differentiated hRGCs were plated at a density of 25,000 cells per well in 96-well clear-bottom, black-walled plates and cultured for 3 days. Subsequently, cells were treated with WAY, DPAT, or WAY with or without PGC1α targeted and non-targeted ASO at specified time points. Caspase activity was assessed using the ApoTox-Glo Triplex assay kit (Promega). For this, 100 μl of Caspase-Glo® 3/7 reagent was added to each well, and the plate was incubated for 30 minutes at room temperature. Luminescence was then recorded to quantify caspase activity. All caspase measurements were normalized to the corresponding control condition.

### Immunofluorescence and confocal imaging

Purified stem cell derived human retinal ganglion cells (hRGCs) were seeded at a density of 30,000 - 40,000 cells per Matrigel-coated coverslip (with 1.5 thickness) and cultured for 3 days. Following treatment with WAY-100635 at defined time points, culture media was removed, and cells were washed with 1× PBS. Cells were fixed with 4% paraformaldehyde for 15 minutes at 37 °C and subsequently permeabilized with 0.5% Triton X-100 in PBS for 5 minutes. Coverslips were then washed three times for 5 minutes each with washing buffer (1% donkey serum and 0.05% Triton X-100 in PBS). Cells were blocked in blocking buffer (5% donkey serum and 0.2% Triton X-100 in PBS) for 1 hour at room temperature. Primary antibodies against Tom20 (Santa Cruz), cAMP (Bio-Techne), 5-HT1A (Abcam), and PGC1α (Abcam) were diluted 1:200 in blocking buffer and applied to coverslips, followed by overnight incubation at 4 °C. The next day, cells were washed three times with washing buffer and incubated for 2 hours at room temperature in the dark with species-specific, fluorophore-conjugated secondary antibodies (1:500 dilution). DAPI (1.43 μM) was added during the second wash to label nuclei. Following final washes, coverslips were mounted onto glass slides using DAKO mounting medium. Fluorescence images were captured using a Zeiss LSM700 confocal microscope equipped with a 63×/1.4 NA oil immersion objective (Fig. 1f, 1j, 2a, 2e, 2h, 2j, 3a, and Supplementary 2a, 5a) and 40x/1.3 NA oil immersion objective (Fig. 1h and Supplementary 1a). Image analysis and quantification were performed using ImageJ software.

### Western blot

Purified hRGCs were seeded at a density of 500,000 cells per well in Matrigel-coated 24-well plates and maintained in culture for 3 days. Cells were then treated with WAY-100635 at indicated time points, with distilled water (dH₂O) serving as the vehicle control, as WAY was dissolved in dH₂O. Following treatment, cells were lysed in 100 μL of M-PER extraction buffer supplemented with 5 mM EDTA and protease inhibitors. Protein concentration was determined using the DC Protein Assay Kit II (Bio-Rad) with bovine serum albumin (BSA) as the standard, and absorbance was read using a microplate reader. Samples were prepared by mixing protein lysates (10 - 20 μg) with 1× Laemmli sample buffer and denaturing at 95 °C for 5 minutes. Proteins were separated on Mini-PROTEAN TGX precast gels (Bio-Rad) using Tris-Glycine-SDS running buffer at 100 V until the dye front reached the bottom of the gel.

Proteins were transferred onto PVDF membranes pre-activated with methanol using Tris-Glycine transfer buffer containing 20% methanol, at 30 V for 2 hours at 4 °C. Membranes were blocked with 5% non-fat dry milk in TBST (Tris-buffered saline with 0.5% Tween-20) for 2 hours at room temperature and then incubated overnight at 4 °C with primary antibodies against Tom70 (Santa Cruz), PGC-1α (Abcam), 5-HT1A (Abcam), and GAPDH (CST), all diluted 1:1000. After washing three times with TBST, membranes were incubated with HRP-conjugated anti-rabbit secondary antibody (CST) diluted 1:10,000 in 5% milk/TBST for 2 hours at room temperature. Following three additional TBST washes, signal was developed using Clarity Max Western ECL substrate (Bio-Rad) and visualized with a Bio-Rad ChemiDoc Gel Imager. Protein bands were quantified by measuring raw integrated density using ImageJ software. Each band was normalized to the corresponding GAPDH (or actin) loading control. Additionally, each treatment condition was normalized to its respective vehicle control. Quantification was based on expected molecular weights as per antibody datasheets and published references.

### Antisense Oligonucleotide (ASO) design and treatment

Non-targeting scrambled control ASOs and PGC-1α-targeting ASOs were designed and synthesized by Integrated DNA Technologies (IDT). All ASOs were modified with IDT’s Affinity Plus chemistry. ASO sequences are listed in Supplemental Table 1. The PGC-1α-targeting ASOs were designed to bind the 3′ untranslated region (3′UTR) of the human PGC-1α mRNA and were applied to H7-WT hRGCs. ASOs (1 µM) were delivered via standard gymnotic uptake (without a transfection reagent). For knock-down analysis by western blots (Fig. 2g), cells were collected after indicated timepoints post first ASO addition and ASOs were replenished every 48 hours for the duration of the experiment. For mitochondrial mass measurements by confocal immunofluorescence (Fig. 2h – k), cells were treated with ASOs (1 µM) for 24 h then treated with WAY (5 µM) by exchanging media containing WAY and ASO for the indicated timepoints.

### JC1 imaging

Purified hRGCs were seeded at a density of 40,000 cells per Matrigel-coated glass-bottom MatTek dish (No. 1.5 thickness) and cultured for 3 days. Cells were then treated with WAY-100635 at designated time points. Following treatment, cells were washed with iNS media and incubated with JC-1 dye (1:100 dilution in iNS; final volume 250 μl) for 30 minutes at 37 °C in a 5% CO₂ incubator. After incubation, the JC-1 media was removed, cells were washed with fresh iNS, and 2 ml of iNS was added before transferring the dish to a live-cell imaging chamber (Tokai Hit) maintained at 37 °C and 5% CO₂. Confocal z-stack images were acquired using a Zeiss LSM700 confocal microscope with a 63×/1.4 NA oil immersion objective. ImageJ was used to generate sum projections of the red (J-aggregates) and green (JC-1 monomers) channels for quantitative analysis. Background red fluorescence, contributed by tdTomato expression was subtracted based on cytoplasmic intensity measurements. Green fluorescence background was measured from cell-free regions and similarly subtracted. Mitochondrial membrane potential was quantified by calculating the red-to-green fluorescence ratio, which was normalized to the corresponding untreated control or average DMSO-treated condition for each cell line.

### Tetramethylrhodamine methyl ester (TMRM) imaging

Purified hRGCs were seeded at a density of 40,000 cells per Matrigel-coated glass-bottom MatTek dish (No. 1.5 thickness) and maintained in culture for 3 days. Cells were treated with WAY-100635 at designated time points. Prior to staining, cells were washed twice with 750 μl of iNS media and then incubated with 250 μl of iNS media containing TMRM (Thermo Fisher) at a final concentration of 100 nM for 30 minutes at 37 °C in a 5% CO₂ incubator. TMRM was freshly prepared from a 100 μM DMSO stock solution by 1:1000 dilution (1.5 μl in 1.5 ml iNS). After incubation, the dye was removed, and cells were washed twice with 750 μl of iNS. Finally,1.5 ml of fresh iNS was added, and dishes were transferred to a live-cell imaging chamber (Tokai Hit) set at 37 °C and 5% CO₂. Confocal z-stack images were acquired using a Zeiss LSM700 confocal microscope equipped with a 63×/1.4 NA oil immersion objective. Image analysis was performed using ImageJ. Sum projections of TMRM fluorescence (red channel) were generated, and background fluorescence contributed by cytoplasmic tdTomato expression in hRGCs was subtracted. Fluorescence intensity was quantified by applying a threshold to define mitochondria-rich regions, and signal was normalized to cell area. All fluorescence measurements were normalized to the average signal of DMSO-treated controls for each experimental condition.

### Multi-electrode Array (MEA)

Stem cell derived hRGCs were seeded at a density of 500,000 cells per well onto MEA plates (CytoView MEA 24-White, Axion Biosystems) and cultured for 11 days in iNS media, with media changes every other day. On day 11, the media was refreshed, and the plate was placed into the Maestro Edge MEA recording system (Axion Biosystems) equipped with environmental control to maintain 37 °C and 5% CO₂. Following a 15-minute equilibration period, spontaneous neuronal activity was recorded for 3 minutes every 3 hours over 24 hours. After this baseline recording phase, WAY-100635 was added directly to the wells at a concentration of 5 μM, and gentle mixing was performed to ensure even distribution. Recordings were then continued every 3 hours for another 24 hours. Data analysis was performed by averaging the mean firing rate from all 16 electrodes per well for each time point. The nine pre-treatment recordings were averaged to determine the baseline activity, while the nine post-treatment recordings were averaged to quantify the effects of WAY treatment.

### Seahorse analysis

Purified hRGCs were seeded onto Matrigel-coated Agilent Seahorse XF96 cell culture microplates at a density of 250,000 cells per well and cultured for 2 days prior to the assay. For hRGC differentiation culture day 2-6 treatment, cells were treated with 5 μM WAY along with the differentiation small molecules. Twenty-four hours before the assay, culture media were replaced with 100 μl iNS containing 5 μM WAY. The day before the assay, Seahorse XF sensor cartridge was hydrated with 200 μl sterile water per well and incubated overnight at 37 °C in a non-CO₂ incubator. On the day of assay, water was replaced with pre-warmed XF calibrant buffer (Agilent), and cartridge was incubated for an additional 45–60 minutes. Seahorse assay medium was prepared using XF DMEM supplemented with 21.25 mM glucose, 0.36 mM sodium pyruvate, and 1.25 mM L-glutamine, with the pH adjusted to 7.38 -7.42 at 37 °C. For different assays, the following compounds were loaded into the cartridge ports: for the Mito Stress Test, 2 μM oligomycin, 2 μM FCCP (optimized from prior titration), and 0.25 μg/ml rotenone + 0.5 μM antimycin A; for the ATP rate assay, 2 μM oligomycin and 0.25 μg/ml rotenone + 0.5 μM antimycin A; and for the glycolytic rate assay, 0.25 μg/ml rotenone + 0.5 μM antimycin A and 17.5 mM 2-deoxy-D-glucose (2-DG). To prepare wells for the assay, 60 μl of iNS was removed from each well and replaced with 140 μl of Seahorse assay medium, followed by another media exchange step using 140 μl fresh assay medium to achieve a final volume of 180 μl per well. Empty wells were filled with 180 μl assay medium. For hRGCs, cell distribution and confluence were recorded using Incucyte S3 (Sartorius) via brightfield and tdTomato fluorescence imaging for normalization. The plate was then incubated at 37 °C in a non-CO₂ incubator for 45 minutes prior to assay. After calibration of the sensor cartridge in the Seahorse XFe96 Analyzer (Agilent) using WAVE software, the cell plate was loaded, and the selected assay was run. Post-assay, image analysis was performed using ImageJ to measure cell area for normalization in hRGCs, while day 2-6 WAY treated or untreated differentiation culture data were normalized using flow cytometry-based cell counts. Data were processed and exported using Seahorse Wave Desktop software and associated Excel macros (Agilent).

### Measurement of hRGC differentiation efficiency

To evaluate hRGC differentiation efficiency, entire wells containing mixed populations of differentiating cells were dissociated into single-cell suspensions using Accutase. Cells were incubated with Accutase for 10-15 minutes at 37 °C, followed by gentle trituration in iNS media to achieve a homogeneous single-cell suspension. The cells were centrifuged, resuspended in fresh iNS media. Single-cell suspensions were analyzed on the Attune NxT flow cytometer (Thermo Fisher). Live cells were first identified using forward and side scatter properties, and doublets were excluded by gating on the singlet population using FSC-A vs. FSC-H plots (**Fig. S8** for gating). Percentage of tdTomato positive hRGCs was then assessed within the singlet population.

### qPCR

hESCs were seeded at a density of 15,000 cells per well in Matrigel-coated 24-well plates and maintained for 7 days. From day 2 to day 6, cells were treated daily with 5 μM WAY-100635, alongside the standard differentiation small molecules. On day 7, the media were aspirated, and cells were incubated with 200 μL Accutase for 10 minutes at 37 °C. The enzymatic reaction was quenched with 400 μL of iNS medium, followed by centrifugation at 150 ×g for 6 minutes. Supernatants were removed, and cell pellets were stored at −20 °C until further processing. Total RNA was extracted using the RNeasy Mini Kit (Qiagen, #74104) according to the manufacturer’s protocol. RNA concentration was quantified using a NanoDrop 2000c spectrophotometer (Thermo Scientific), and 6 μL of RNA was used for cDNA synthesis using the Abcam cDNA Synthesis Kit (#G492). Quantitative PCR (qPCR) was performed using BlasTaq™ qPCR MasterMix in a 20 μL reaction containing 100 ng of total cDNA per well, on a QuantStudio 6 Flex Real-Time PCR system (Applied Biosystems). Primers used for gene expression analysis are listed in Supplemental Table 1. GAPDH was used as a housekeeping gene for normalization. ΔCt values were calculated relative to the housekeeping gene, and ΔΔCt values were determined relative to the mean ΔCt of control (untreated) samples. Each condition was measured in triplicate (technical replicates), with 3-4 independent biological replicates.

### Cell cycle stage assessment

Cell cycle distribution was assessed by measuring DNA content using propidium iodide (PI) staining, following an established protocol^51^. Cells were dissociated using Accutase, pelleted, and fixed overnight at –20 °C in 70% ice-cold ethanol. Following fixation, cells were washed twice with ice-cold PBS and resuspended in PBS containing 20 μg/ml PI, 0.1% Triton X-100, and 100 μg/ml RNase A. The suspension was incubated at room temperature for at least 30 minutes to allow sufficient DNA staining. Flow cytometric analysis was performed using the Attune NxT flow cytometer (Thermo Fisher). Doublets were excluded by gating on the singlet population (FSC-A vs. FSC-H), and PI fluorescence was measured to quantify DNA content. Data were analyzed using ModFit LT software (Verity Software House), which modeled the PI fluorescence distribution to determine the proportion of cells in G0/G1, S, and G2/M phases based on DNA content. Results were visualized as histograms showing phase-specific peaks with corresponding percentage values.

### Electron microscopy

hESCs were seeded at a density of approximately 350,000 cells per well in 6-well Matrigel-coated plates and maintained for 7 days. From day 2 to day 6, cells were treated with 5 μM WAY-100635 along with the standard differentiation small molecules. In another 6-well plate, hESCs were seeded following routine passaging procedures using Gentle Cell Dissociation Reagent when cultures reached 70–80% confluency. These cells were maintained until they reached 80-90% confluency and were then used as pure stem cell samples for EM without any differentiation. On day 7, media were aspirated and ∼1 million cells per condition were collected and pelleted by centrifugation at 1000 × g for 10 minutes at 4 °C. Cell pellets were fixed overnight at 4 °C in 1 mL of 0.1 M cacodylate buffer (pH 7.4) containing 2.5% glutaraldehyde and 2% paraformaldehyde (Electron Microscopy Sciences, Hatfield, PA, USA). Pellets were washed three times with 4 mL of 0.1 M cacodylate buffer, then subjected to secondary fixation with 4 mL of 0.1 M cacodylate buffer containing 1% osmium tetroxide (OsO₄) and 1.5% potassium ferrocyanide for 1 hour at room temperature. Samples were dehydrated through a graded ethanol series: 30% ethanol for 10 minutes, 50% for 30 minutes, 70% for 30 minutes, followed by two washes in 95% ethanol for 15 minutes each, and two washes in 100% ethanol for 30 minutes each. Infiltration was performed on a rotator at 60 rpm with SPURR resin (Electron Microscopy Sciences) mixed in increasing concentrations: 1:2 resin: ethanol for 16 hours, 1:1 for 24 hours, 3:1 for 3 hours, and 100% SPURR for 6 hours. Pellets were embedded in BEEM capsules filled with 400 μL of fresh SPURR resin and polymerized by incubating at 70 °C for 12 hours. Ultrathin sections (70 nm) were cut using a Leica ultramicrotome with a Diatome diamond knife and mounted on 200-mesh nickel grids. Grids were dried in a vacuum desiccator (SP Bel-Art, Wayne, NJ, USA) for 1 hour before imaging. Electron micrographs were acquired using a Tecnai Spirit BioTwin transmission electron microscope (FEI, Hillsboro, OR, USA) operating at 80 kV, equipped with an AMT NanoSprint 6 CMOS camera (AMT, Woburn, MA, USA).

### Animals

All animal experiments were approved by the Institutional Animal Care and Use Committee at the Indiana University School of Medicine and conducted in accordance with the ARVO Statement for the Use of Animals in Ophthalmic and Vision Research and the ARRIVE guidelines. Wild-type C57BL/6J male mice (2-3 months old) were obtained from Jackson Laboratory (Bar Harbor, ME, USA) and housed under standard conditions at the Laboratory Animal Resource Center, Indiana University School of Medicine. In female mice, the estrous cycle-with its fluctuating levels of estrogen and progesterone, influences both neuronal injury and recovery processes. These hormonal variations modulate neuronal excitability, neurovascular dynamics, and the expression of neurotrophic factors^107,108^. Estrogen, in particular, plays a critical role in regulating mitochondrial function, thereby impacting cellular energy metabolism and neuronal resilience^109^. Given these influences, it is important to consider the stage of the estrous cycle when assessing outcomes in studies involving neuronal injury and regeneration. To avoid these complications, we have studied neuronal injury only in male mice. For the optic nerve crush (ONC) model, mice were divided into three groups (n = 3-6 per group): naïve, ONC, and ONC+WAY. All animals were acclimatized for one week prior to experiments. The ONC+WAY group received daily intraperitoneal injections of WAY-100635 (5 mg/kg body weight) for 5 days before ONC surgery. ONC was performed on day 0 in both the ONC and ONC+WAY groups, and WAY treatment was continued post-ONC for up to day before sacrifice, depending on the experimental timeline. In an alternate cohort, post-treatment with WAY began 4 hours after ONC and continued daily until day before sacrifice. At the end of each experiment, animals were euthanized, and eyes, optic nerves, kidneys, and livers were collected. Body weight was recorded at 5 days prior to ONC, on the day of ONC, and at the time of sacrifice.

For the elevated intraocular pressure (IOP) model using microbead injection, mice were assigned to three groups (n = 3 per group): PBS injection (sham), microbead-injected (MB), and microbead-injected plus WAY treatment (MB+WAY). All animals were acclimatized for one week. The MB+WAY group received daily intraperitoneal injections of WAY (5 mg/kg body weight) for 6 days prior to microbead injection, which was performed on day 0 in both the MB and MB+WAY groups. WAY treatment continued daily post-injection. At the end of the study, animals were euthanized, and eyes were harvested for further analysis.

### Optic Nerve Crush mouse model

Mice were anesthetized using 1.5-2% isoflurane (Akorn, Illinois, USA), and 0.5% proparacaine hydrochloride ophthalmic drops (Bausch + Lomb, Laval, Canada) were administered to both eyes for local anesthesia. Phenylephrine Hydrochloride ophthalmic Solution (Akorn, Illinois, USA) was also administered to dilate the pupils. A small incision was made in the upper conjunctiva, and the eyeball was gently retracted outward using fine forceps. A second pair of forceps was used to carefully open the connective tissue surrounding the optic nerve. Under a dissecting microscope, the optic nerve was located and crushed approximately 1 mm behind the globe using calibrated forceps for a duration of 5 seconds. Following the optic nerve crush (ONC), the eye was gently repositioned into the orbit, and the mouse’s head was released. To prevent infection and ensure eye lubrication during recovery, erythromycin ophthalmic ointment USP, 0.5% (Bausch + Lomb, Laval, Canada) was applied. Animals were monitored continuously until full recovery from anesthesia was confirmed.

### Bead injected high IOP mouse model

Mice were anesthetized with 1.5-2% isoflurane, and 0.5% proparacaine hydrochloride eye drops were applied for topical anesthesia. Phenylephrine Hydrochloride ophthalmic Solution was also administered to dilate the pupils. Under general anesthesia, microbeads were injected unilaterally into the anterior chamber of the mouse eye following established methods^110^. Briefly, <50 μm glass cannulas connected to a Hamilton syringe were used for injection. Microbeads were sterilized by placing them in 100% ethanol in 0.5 mL Eppendorf tubes, followed by centrifugation, resuspension in ethanol, and repeating this wash step twice. They were then washed and resuspended in sterile phosphate-buffered saline (PBS), and the final bead pellet was aspirated directly into a glass micropipette. For injection, a glass cannula was loaded with a mixture of 1 μm and 6 μm microbeads combined with the viscoelastic solution Healon (10 mg/mL sodium hyaluronate, Johnson & Johnson), which ensured maximum retention of the beads after cannula withdrawal. The left eye was gently proptosed and the cannula was inserted into the inferior region of the anterior chamber. The bead solution was slowly injected over a 45-second period, and the cannula was left in place for 2 minutes post-injection to minimize efflux. Mice were monitored carefully post-procedure until they fully regained consciousness.

### Intraocular pressure measurement

Intraocular pressure (IOP) was measured using the TonoLab rebound tonometer (TioLat, Inc., Helsinki, Finland). For each eye, five consecutive measurements with optimal quality grading were recorded and averaged to obtain a final IOP value. Baseline IOP was measured prior to microbead injection, and subsequent measurements were taken at designated time points following the injection to monitor changes in IOP over time.

### Retinal flat mounting and RGC count

Retinal tissues were collected and immersed in ice-cold PBS before being fixed in 4% paraformaldehyde. Fixed tissues were then incubated five overnights with either guinea pig or rabbit anti-RBPMS antibody (catalogue no. 1832-RBPMS or 1830-RBPMS, PhosphoSolutions, Aurora, CO) at a dilution of 1:200. To detect the primary antibody, an Alexa Fluor 647-labeled goat anti-guinea pig secondary antibody (catalogue no. A21450, Invitrogen, Carlsbad, CA) or Donkey anti-rabbit Alexa Fluor 647 (catalogue no. A31573, Invitrogen, Carlsbad, CA) was used. Nuclei were counterstained with DAPI during the final wash step. Eye cups were cut in a flower-petal shape and flat-mounted on slides using 0.45 μm black mixed cellulose ester (MCE) membrane filters. Coverslips were applied using Aqua-Poly/Mount mounting medium. Confocal immunofluorescence imaging was performed with a Zeiss LSM700 microscope using a 40x/1.3 NA oil objective. For each retina, eight z-stacked images were acquired from four quadrants, sampling both central and peripheral regions. Manual quantification of RBPMS-positive RGCs was performed on maximum intensity projections of the images.

### Optic nerve cryosection preparation and staining

Cryosections are prepared following published methods^111^ but with modifications. Optic nerves were fixed in 4% paraformaldehyde prepared in 0.1 M phosphate buffer (PB; NaH₂PO₄ and Na₂HPO₄ in Milli-Q water, pH 7.4) for 6 hours at room temperature (RT). Following fixation, tissues were transferred to a 5% sucrose solution in phosphate buffer for 30 minutes, then sequentially incubated in increasing concentrations of sucrose (10%, 15%, and 20%) in 0.1 M PBS. For cryopreservation, tissues were incubated overnight at 4°C in a mixture of 20% sucrose and OCT compound (Sakura Finetek) at a 2:1 ratio. The next day, tissues were oriented in Tissue-Tek cryo molds (Sakura Finetek) and snap-frozen in isopentane cooled with liquid nitrogen. Frozen tissues were sectioned using a Leica CM1850 cryostat (Leica Microsystems) at a thickness of 14 μm and mounted on Super Frost Plus microscope slides. After drying, the slides were stored at -80°C until use. Prior to staining, slides were warmed in a 37°C incubator and tissue sections were outlined with an ImmEdge hydrophobic barrier pen (Vector Laboratories).

For staining, sections were first washed in Tris-buffered saline (TBS, pH 7.4) for 5 minutes, followed by pre-incubation in 100% methanol for 10 minutes. Blocking was performed using a solution of 10% donkey serum in TBS. Sections were then incubated overnight at 4°C with a primary antibody against growth-associated protein 43 (GAP43) (1:1,000; custom-made in sheep by Dr. Benowitz)^112–114^, diluted in solution A (TBS2T: 300 mM NaCl, 0.1% Tween 20, pH 7.4) containing 2% BSA and 10% donkey serum. The following day, sections were rinsed in TBS2T for 1 hour at 4°C, followed by a 1-hour rinse in solution A at room temperature, and then a final rinse in TBS2T for 1 hour at room temperature. Sections were subsequently incubated for 2 hours at room temperature with an AlexaFluor-488-conjugated donkey anti-sheep IgG secondary antibody (1:500; Thermo Fisher Scientific, #A11015), followed by washes in TBS2T (2 × 5 min) and TBS (5 min) at room temperature. Sections were mounted using one drop of DAKO mounting medium, coverslip, sealed with nail polish (Electron Microscopy Sciences), and allowed to dry overnight at room temperature before imaging. Confocal stitched immunofluorescence images were taken using Zeiss LSM700 microscope or Nikon AX R confocal microscopes. For quantification, GAP43-positive axons crossing a virtual line located at 0.5 mm distal to the optic nerve crush site were counted. Four sections per nerve, spaced 28 μm apart, were analyzed. The maximum section width was used to estimate the cross-sectional area at 0.5 mm, and this, along with section thickness and axon counts, was used to estimate the total number of regenerating axons at that distance following published method by Benowitz lab^60^.

### Retinal cross-sections and immunostaining

Mice were euthanized by CO₂ inhalation followed by cervical dislocation. Eyes were enucleated and briefly rinsed with PBS. Two small punctures were made at the junction of the cornea and sclera using a needle, and the eyes were fixed in 4% paraformaldehyde (PFA) in PBS for 15 minutes. The cornea was removed, followed by extraction of the lens and vitreous body. Retina was isolated. The retina was further fixed in 4% PFA for 60 minutes at room temperature. After fixation, eyecups were washed three times in PBS (5 minutes each) and cryoprotected by immersion in a graded sucrose series in PBS (5%, 10%, 15%, and 20% for 30 minutes each at room temperature), followed by overnight incubation at 4°C in a 2:1 mixture of sucrose-PBS and OCT compound. The next day, the tissues were embedded in OCT (Tissue-Tek) and snap-frozen in liquid nitrogen vapor. Eyecups were cryosectioned at 10 μm thickness using a Leica CM3050S cryostat and stored at −80°C until use.

For immunohistochemistry, cryosections were thawed and air-dried at 37°C for 40 minutes. Sections were washed three times with 0.1% PBST (PBS + 0.1% Triton X-100), then blocked in PBST containing 5% goat serum for 16-24 hours at 4°C. Following blocking, sections were rinsed once and washed three times (5 minutes each) with PBST, then incubated overnight at 4°C with primary antibodies diluted in PBST. The next day, sections were again rinsed once and washed three times with PBST before incubation with secondary antibodies and DAPI in PBST for 1 hour at 4°C. After final washes (three times for 5 minutes each), sections were mounted using DAKO mounting medium and coverslips. Confocal immunofluorescence imaging was performed with a Zeiss LSM700 microscope using a 40x/1.3 NA oil objective.

### Photopic negative response (PhNR) recording by flash electroretinogram (ERG)

RGC function was assessed using the Celeris system (Diagnosys, Inc., Lowell, MA) in conjunction with Espion software (Diagnosys, Inc.). Electroretinograms (ERGs) were recorded 12 days following optic nerve crush (ONC). Mice were dark-adapted overnight, and all procedures were performed under dim red light (660 nm). Animals were anesthetized with isoflurane and positioned on a heated platform maintained at 37 °C. Pupils were dilated using 1% tropicamide (Somerset Therapeutics, FL), and topical anesthesia was applied using 0.5% proparacaine hydrochloride (Bausch + Lomb, NJ). Corneal hydration was maintained using Systane (Alcon, TX). A contact lens-type probe with an integrated electrode and stimulator, featuring a pinhole aperture, was placed on the cornea and aligned with the center of the pupil. The reference electrode was placed on the contralateral eye. PhNR recordings were obtained using 100 consecutive white light flashes (20.0 cd·s/m²) on a rod-saturating green background (40 cd·s/m²) to elicit RGC-specific responses. The raw signal was filtered with a 0.125-40 Hz bandpass filter to minimize oscillatory artifacts. For each animal, the probe was positioned at three different angles to ensure consistent recording, and the PhNR amplitudes were averaged. PhNR amplitude was defined as the voltage from the baseline to the trough immediately following the b-wave, and implicit time (time to trough) was also recorded. After the procedure, animals were placed back in their home cages on heating pads covered with drapes. Cages were positioned such that only half rested on the heating pad (low temperature setting), allowing the mice to self-regulate body heat. Once fully ambulatory (∼10-15 minutes), animals were returned to the animal facility.

### Optokinetic response (OKR) test

Following unilateral microbead (MB) left eye injection in mice pretreated with WAY-100635 (5 mg/kg body weight) for 6 days, daily post-treatment with WAY was continued for designated durations, and optokinetic response (OKR) testing was performed on Days 7 and 30 post-MB injections. The OKR test was used to assess visual function, specifically visual acuity (VA) and contrast sensitivity (CS). Mice were placed on a platform surrounded by four computer screens displaying moving sinusoidal gratings (12°/s) in either clockwise or counterclockwise directions. For VA testing, the gratings were presented at increasing spatial frequencies while maintaining fixed speed and contrast. Head-tracking movements in response to the gratings were recorded, with higher frequencies indicating better visual acuity. For CS testing, gratings were presented at a fixed spatial frequency and speed, but with decreasing contrast. CS was quantified as the lowest contrast eliciting a tracking response, reported as Michelson contrast: (max − min) / (max + min), where “max” and “min” represent the maximum and minimum contrast levels to which the mouse responded^115,116^. VA and CS values were recorded separately for clockwise and counterclockwise stimulus directions, as well as combined to obtain an averaged measure of visual performance.

### Histology

The kidney and liver tissues from the naïve and WAY-treated mice were collected and fixed in 10% formaldehyde and processed for paraffin sectioning. Paraffin-embedded sections were stained with hematoxylin and eosin to study the histological changes under a light microscope.

### Mass spectroscopic analysis

To evaluate whether WAY-100635 crosses the blood-retinal barrier, C57BL/6J mice were intraperitoneally injected with WAY-100635 (100 μL in PBS) at a dose of 30 mg/kg body weight. Mice were sacrificed at 4- and 24-hours post-injection, and both blood and eye tissues were collected for analysis. For plasma collection, 75 μL of blood was drawn into heparinized microcapillary tubes and centrifuged at 5,000 rpm for 5 minutes. The separated plasma was transferred to fresh tubes and stored at −80°C. For retinal analysis, the eyes were enucleated, and the retinas were rapidly dissected, flash frozen in liquid nitrogen, and stored at −80°C until further use. Quantification of WAY-100635 in both plasma and retinal tissues was performed using mass spectrometry.

### High-Resolution diffusion MRI and tractography of mouse brains

High-resolution ex vivo MRI was performed using a 9.4 Tesla Bruker 94/30 system (Billerica, MA, USA) with a 30 cm bore and a maximum gradient strength of 660 mT/m on each axis. Diffusion-weighted images (DWI) were acquired using a multi-shot 2D echo-planar imaging (EPI) sequence with the following parameters: matrix size = 100 × 86, field of view = 15 × 12.9 mm², in-plane resolution = 150 μm, slice thickness = 0.3 mm, echo time (TE) = 27 ms, repetition time (TR) = 3500 ms, and b-values of 600, 1200, 1800, and 3000 s/mm², along with four non-diffusion-weighted (b0) volumes. The acquired DWI data were denoised using the Marchenko-Pastur principal component analysis (MP-PCA) method^117^, corrected for Gibbs artifacts via a local subvoxel shift algorithm^118^, and registered nonlinearly to structural T2-weighted images using symmetric normalization (SyN) to correct for EPI distortions, motion, and eddy currents^119^. Bias field correction was applied using the N4 algorithm implemented in ANTs^120^. Diffusion modeling included fitting the diffusion tensor imaging (DTI), diffusion tensor imaging (DKI), and neurite orientation dispersion and density imaging (NODDI) models to extract key diffusion metrics such as fractional anisotropy (FA), mean diffusivity (MD)^72^, kurtosis fractional anisotropy (KFA), mean kurtosis (MK)^73^, neurite density index (NDI), and orientation dispersion index (ODI)^74^. Brain masks were manually validated, and average B0 images were registered to a T2-weighted brain atlas using diffeomorphic registration. Atlas labels were then transformed back into subject space for anatomical reference. For tractography, tissue response functions were automatically derived from six target regions: optic tract (OT), lateral geniculate nucleus (LGN), and visual cortex (VC) bilaterally using MRtrix3. Fiber orientation distributions (FODs)^121^ were computed via constrained spherical deconvolution, followed by probabilistic tractography (10 million streamlines, threshold 0.17)^122^ and SIFT filtering to reduce reconstruction bias^123^. Key tracts between the optic tract, LGN, and visual cortex were extracted bilaterally to evaluate visual pathway connectivity.

### Mouse RGC Purification

Mouse eyes were dissected, and the retinae were isolated and enzymatically dissociated in 500 μL of 5 U/mL Dispase (stored at -20°C) following mechanical trituration. After a 25-minute incubation at 37°C, the dissociated cells were blocked in HBSS containing 2% BSA, counted, and centrifuged at 150 g for 6 minutes. If no pellet was visible, a second centrifugation at 800 g for 6 minutes was performed. Cells were then incubated with anti-CD90.2 (Thy1.2) magnetic microbeads (3 μL beads in 90 μL MACS BSA per 10 million cells) for 30 minutes, washed, and subjected to magnetic-activated cell sorting (MACS) using sequential small columns prewashed with MACS BSA. After three washes per column and elution, the purified RGCs were collected by centrifugation. For Western blotting, the cell pellet was lysed in 100 μL of MPER buffer supplemented with EDTA and protease inhibitors. For flow cytometry, cells were centrifuged at 800 rpm for 7 minutes, resuspended in blocking buffer, fixed and permeabilized with chilled acetone (1:1), and incubated at -20°C for 10 minutes. After washing and pelleting, cells were incubated with primary antibodies (1:100) Tuj1 for 25 minutes at room temperature, washed, and incubated with secondary Alexa Fluor 488 antibody, (1:1000) for 25 minutes. Labeled cells were washed, resuspended in PBS, and analyzed by flow cytometry.

### Statistics and reproducibility

Samples treated with WAY-100635 or 8-OH-DPAT at different time points were considered independent biological replicates. For comparisons between two independent datasets, unpaired two-tailed Student’s t-tests were performed; each data point represents an individual sample (Fig. 1b, 3d, 3f, 5d, 5g–j, 5q, 7c, S2b, d, S6b). For comparisons involving three or more groups with a single independent variable, one-way ANOVA was used (Fig. 1c-d, 1g, 1i, 1k, 2b, 2d, 2f, 2i, 2k-l, 3b, 3i-j, 5c, S1b, 3c). When data involved two independent variables, two-way ANOVA was performed to assess main effects and interactions (Figs. 3h, 4b-c, 4e-g, 6c, 6e, 6g, 6k, 7g-i, S4b). Post-hoc comparisons were corrected using Dunnett’s test when comparing to a control group, or Tukey’s test when comparing among all groups. Graphs were generated using GraphPad Prism 10.0, and figures were assembled in Adobe Illustrator.

## Supplementary Figures

**Fig. S1.**
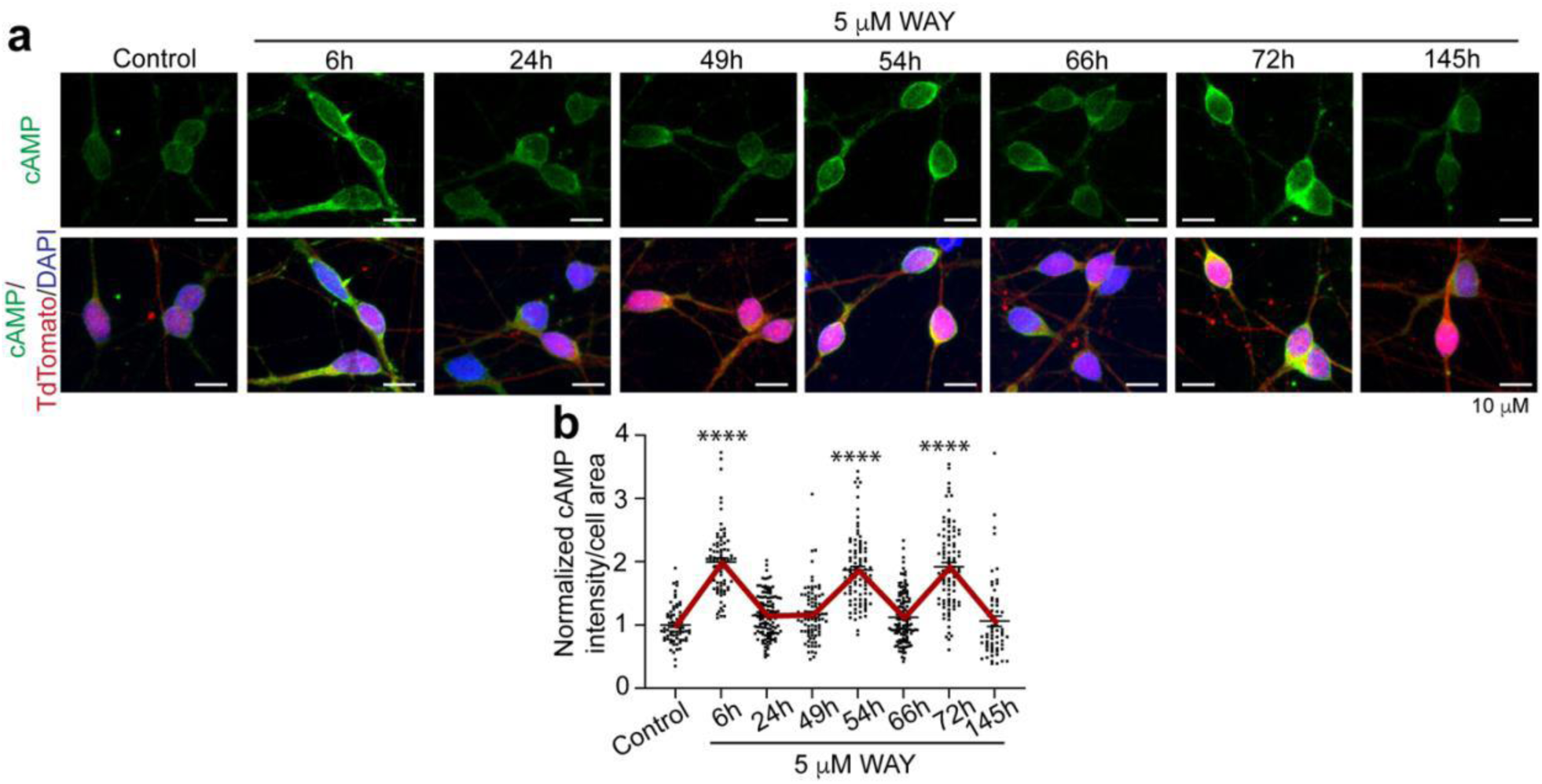
Prolonged WAY-100635 treatment elicits repeated, non-desensitizing cAMP oscillations over six days. **(a)** Representative confocal images (40x/1.3NA, oil) (max projection of z-stacks) of H7-hRGC^WT^ expressing tdTomato (red), and stained for nucleus (DAPI, blue), mitochondria (Tom20, green) at the indicated times after 5 µM WAY. Following the initial treatment, fresh WAY was added at every 24 h. (**b**) Quantification of Tom20 fluorescence intensity per unit cell area from sum projections of confocal z-stacks, normalized to control. Error bars are SEM. One-way ANOVA with Dunnett’s, **** p < 0.0001. n = 53 -129 cells.

**Fig. S2.**
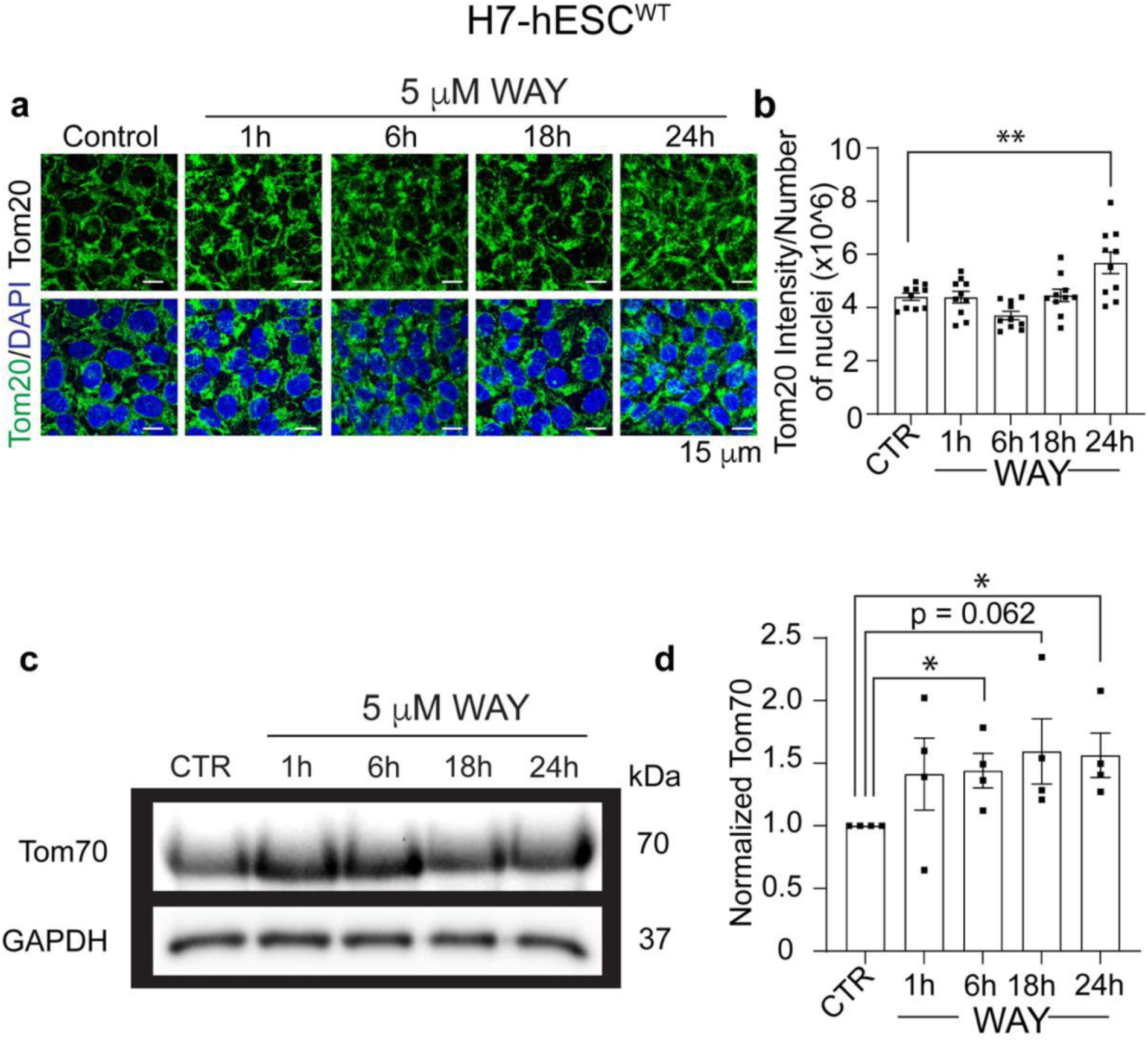
Treatment of hESCs with WAY 100635 (5 µM) led to a significant increase in mitochondrial content. (**a**) Representative confocal images of H7-hESCs^WT^ with Tom20 and DAPI after WAY treatment at indicated timepoints. (**b**) Quantification of Tom20 intensity per image from sum projections of confocal z-stacks at indicated time points, normalized with nuclei number and then control (n = 10 images). **(c**) Representative western blot images of Tom70 with H7-hESCs^WT^ treated with WAY for indicated timepoints. (**d**) Protein band intensity normalized with-respect-to corresponding GAPDH and successively with the control (CTR) to identify change over time (n=4). Error bars are SEM. Unpaired two-tailed t-test, *, p < 0.05, **, p < 0.008.

**Fig. S3.**
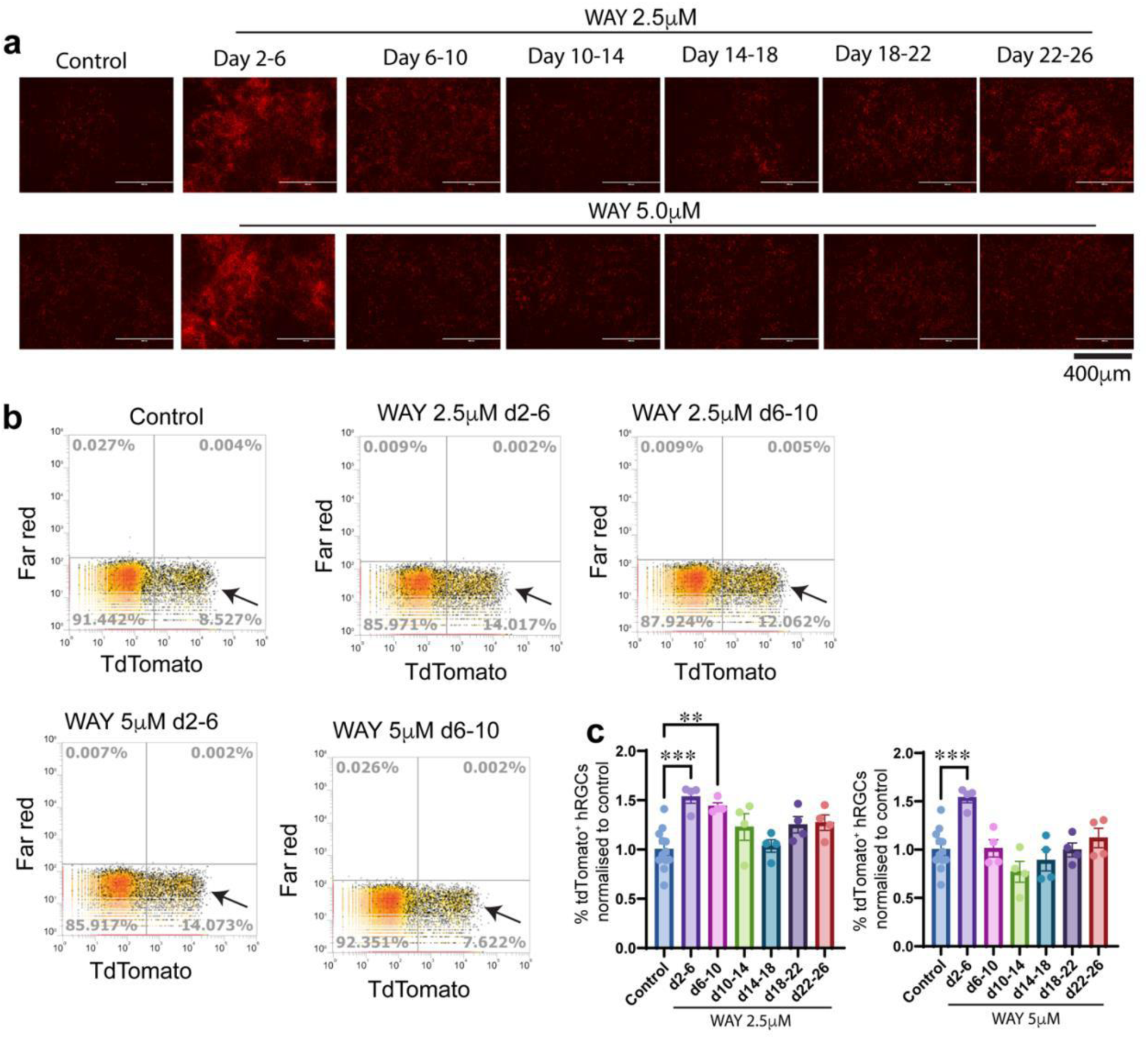
A single WAY-100635 exposure during early in differentiation significantly enhances hRGC yield. **(a**) Live cell fluorescent imaging of TdTomato expressing hRGCs on day 32 under control (untreated) or WAY treatment (2.5 mM and 5 mM) during different time windows (Day 2 - 6, 6 - 10, 10 - 14, 14 - 18, 18 - 22, 22 - 26). (**b)** Cells were collected and analyzed by flow cytometry on day 32. Representative flow cytometry plots with varying population of hRGCs (indicated by arrow) are shown for both concentrations of WAY treatment during days 2 - 6 and days 6 - 10. **(c)** Quantification of TdTomato positive hRGC percentages on day 32 differentiation cultures normalized to control from above treatment conditions. One-way ANOVA with Dunnet’s, *** p < 0.001, ****, P < 0.0001, n = 4 - 12 biological repeats.

**Fig. S4.**
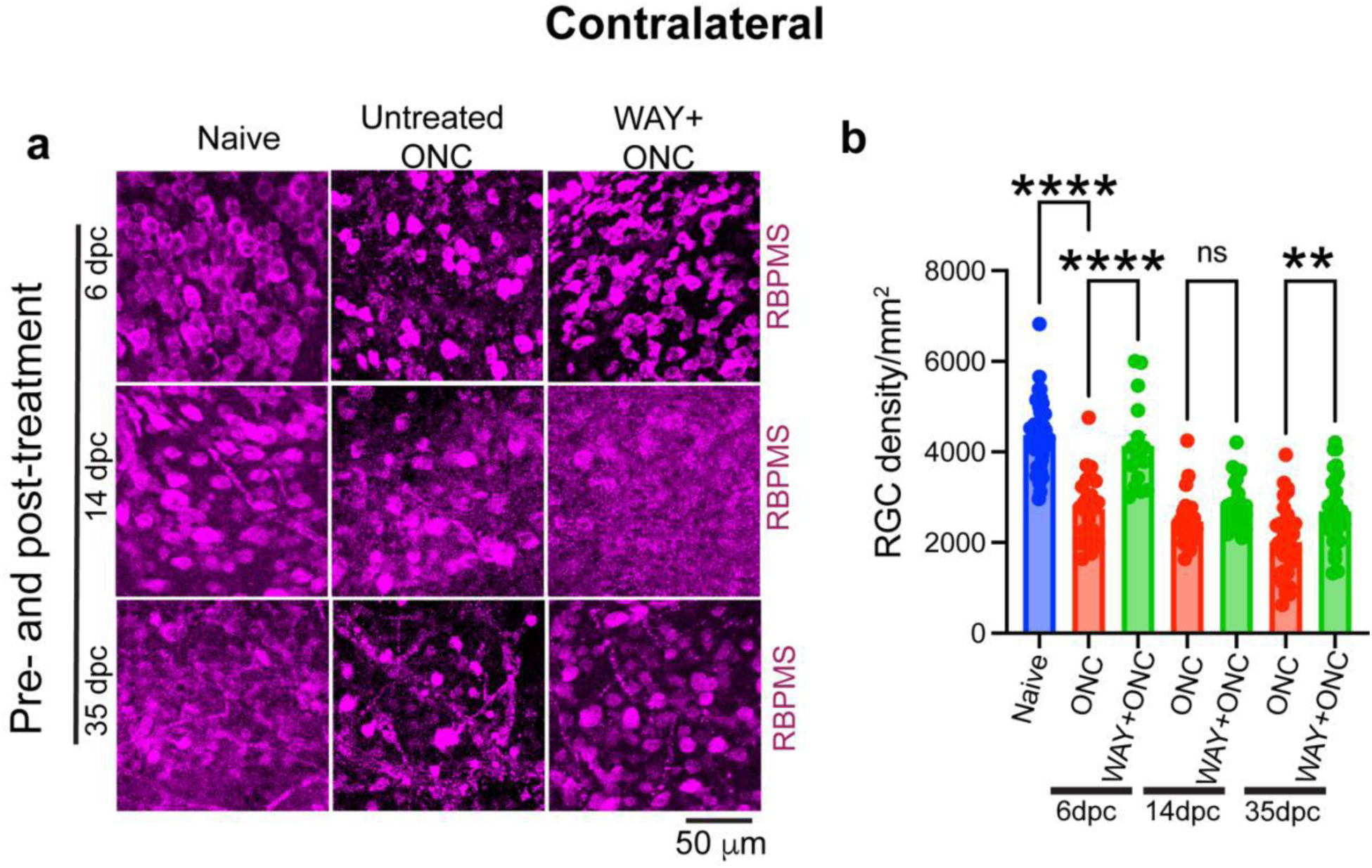
Systemic WAY-100635 preserves RGCs in the contralateral retina. (**a**) Representative confocal micrographs (40 ×/1.3 NA) of retinal flat mounts from mice given daily intraperitoneal WAY (5 mg/kg) starting 5 d before optic nerve crush and continuing for 6, 14, or 35 d post-injury. (**b**) RGC density (cells/mm^2^) from peripheral and central retina under the indicated conditions. Data are mean ± SEM; two-way ANOVA with Tukey’s multiple-comparison test, ****p < 0.0001, **p < 0.01; n = 16 - 48 images from 3 - 6 mice/group. Naïve-eye images and counts are the same as those in Fig. 6b - c.

**Fig. S5.**
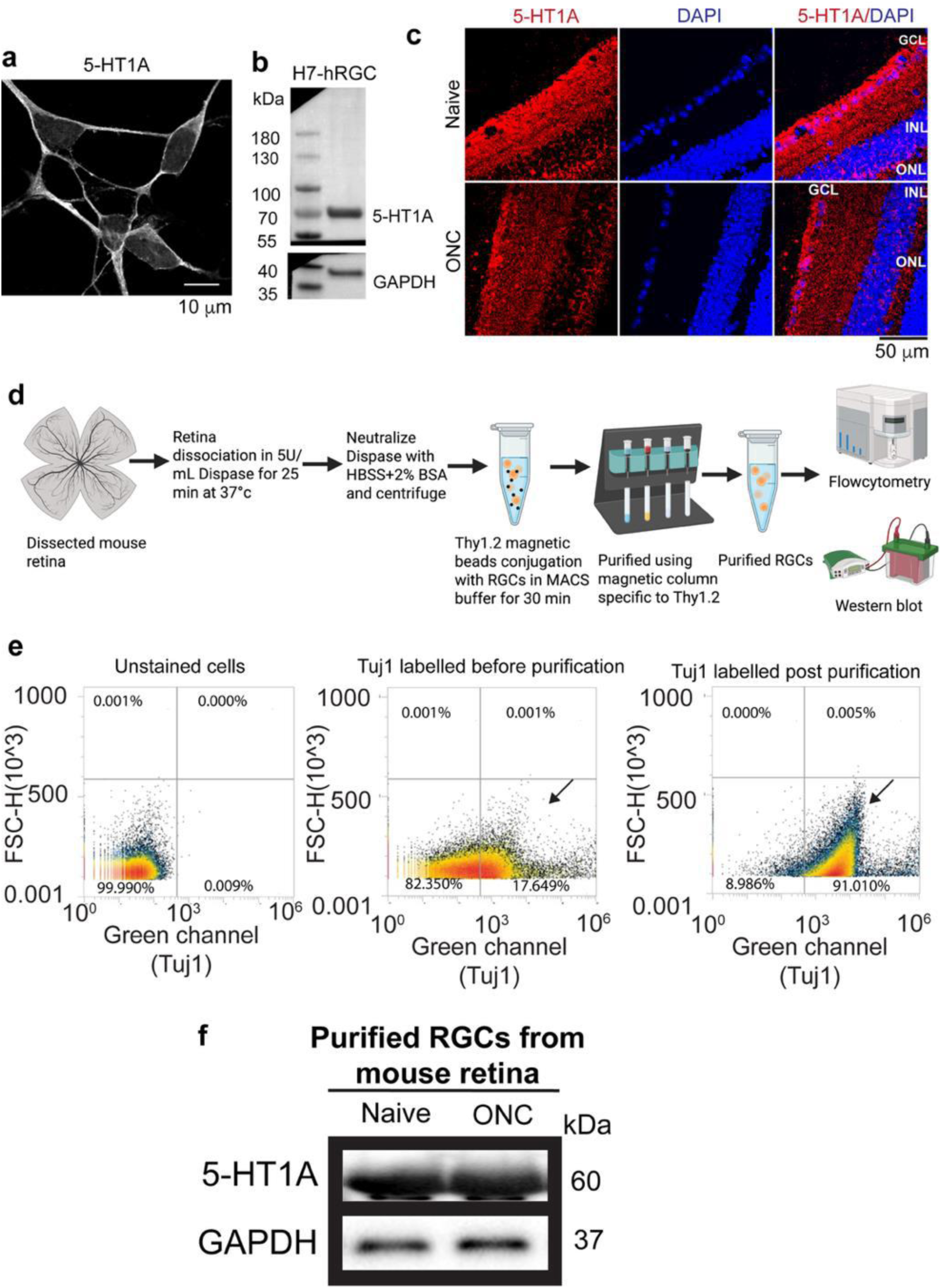
**5-HT1A receptor is abundantly expressed in RGCs and remains stable under optic nerve injury**. (**a**) Representative confocal micrograph of H7-hRGCs immunostained for 5-HT1A; signal is concentrated along the plasma membrane. (**b**) Immunoblot of hRGC lysates confirms 5-HT1A protein; GAPDH serves as a loading control. (**c**) Confocal immunohistochemistry micrographs of mouse retinal cryosections reveals strong 5-HT1A expression in the ganglion-cell layer (GCL - red; nucleus - DAPI) in naïve eyes and remains stable in the remaining ganglion cells 25 d after ONC. (**d**) Schematic of RGC isolation from mouse retinas using CD90.2 (Thy1.2) magnetic beads under chilled conditions, followed by flow cytometry and western blot analysis. (**e**) Flow cytometry of purified cells immunolabeled for the RGC marker Tuj1 (β-tubulin III) demonstrates more than 90 % enrichment. (**f**) Western blot of purified mice RGCs (pooled from three retinas) shows robust 5-HT1A expression from naïve retinas and 24 h post-ONC.

**Fig. S6.**
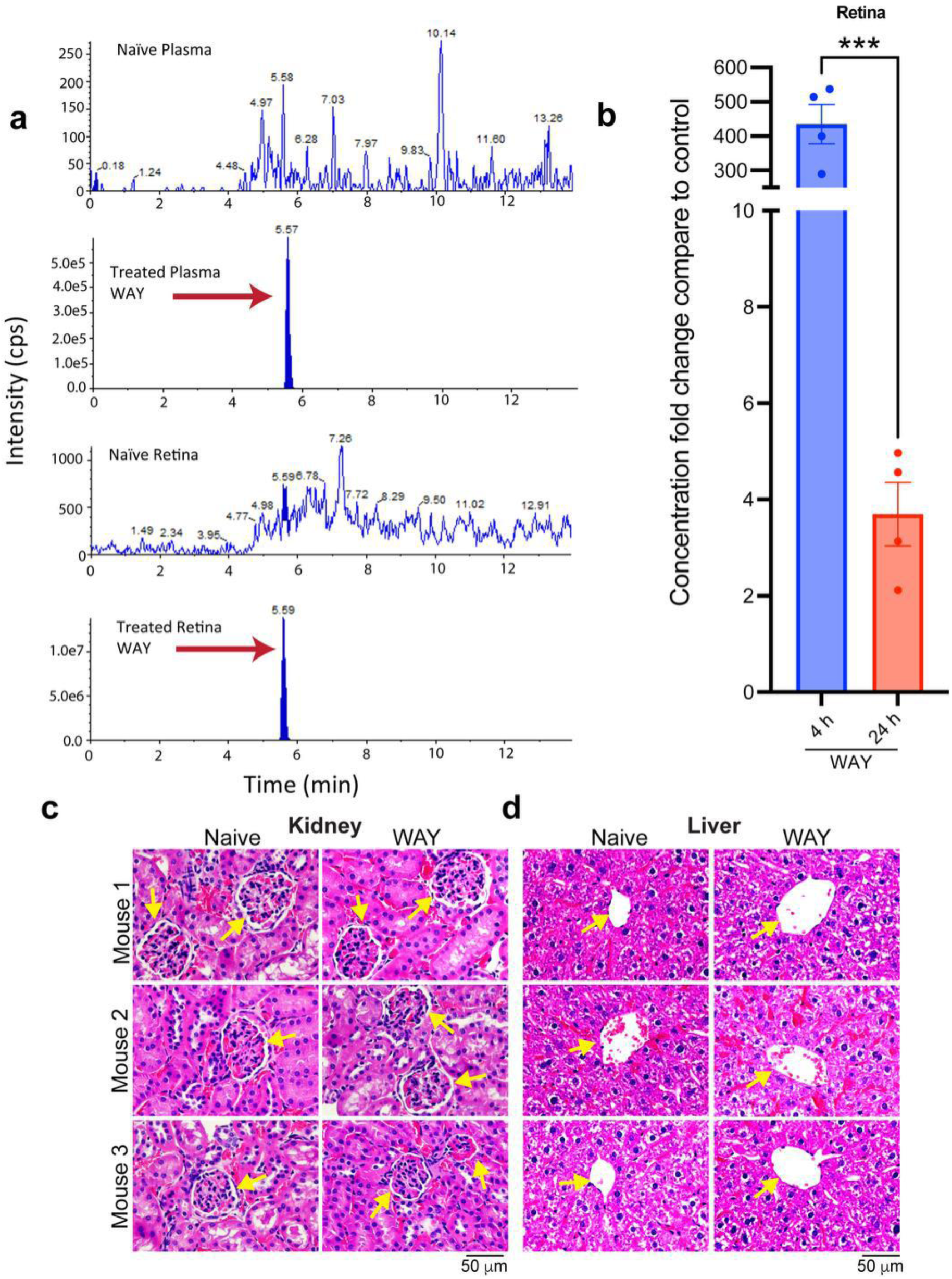
**Systemic WAY-100635 penetrates the retina without producing detectable systemic toxicity**. (**a**) Liquid chromatography-mass spectrometry (LC-MS) chromatograms of plasma and retinal extracts from C57BL/6J mice collected 4 h and 24 h after a single intraperitoneal dose of WAY (30 mg/kg). The blue peak (retention time = 5.5 min) identifies the parent compound in 4 h samples; no corresponding peak is present in naïve (untreated) controls. Trace shown is representative of four biological replicates. (**b**) Retinal WAY concentration (ng/g, mean ± SEM) peaks at 4 h (2,006 ± 264) and falls sharply by 24 h (17 ± 3). Unpaired two-tailed Student’s t-test, ***p < 0.001; n = 4 samples per time point. (**c - d**) Chronic exposure is well tolerated. Hematoxylin-and-eosin sections (40 × / 0.6 NA) of kidney (**c**) and liver (**d**) from naïve mice and mice given daily intraperitoneal WAY (5 mg/kg) for 30 d show normal histoarchitecture with no evidence of inflammation, necrosis, or other pathology. Data are representative of three independent experiments.

**Fig. S7.**
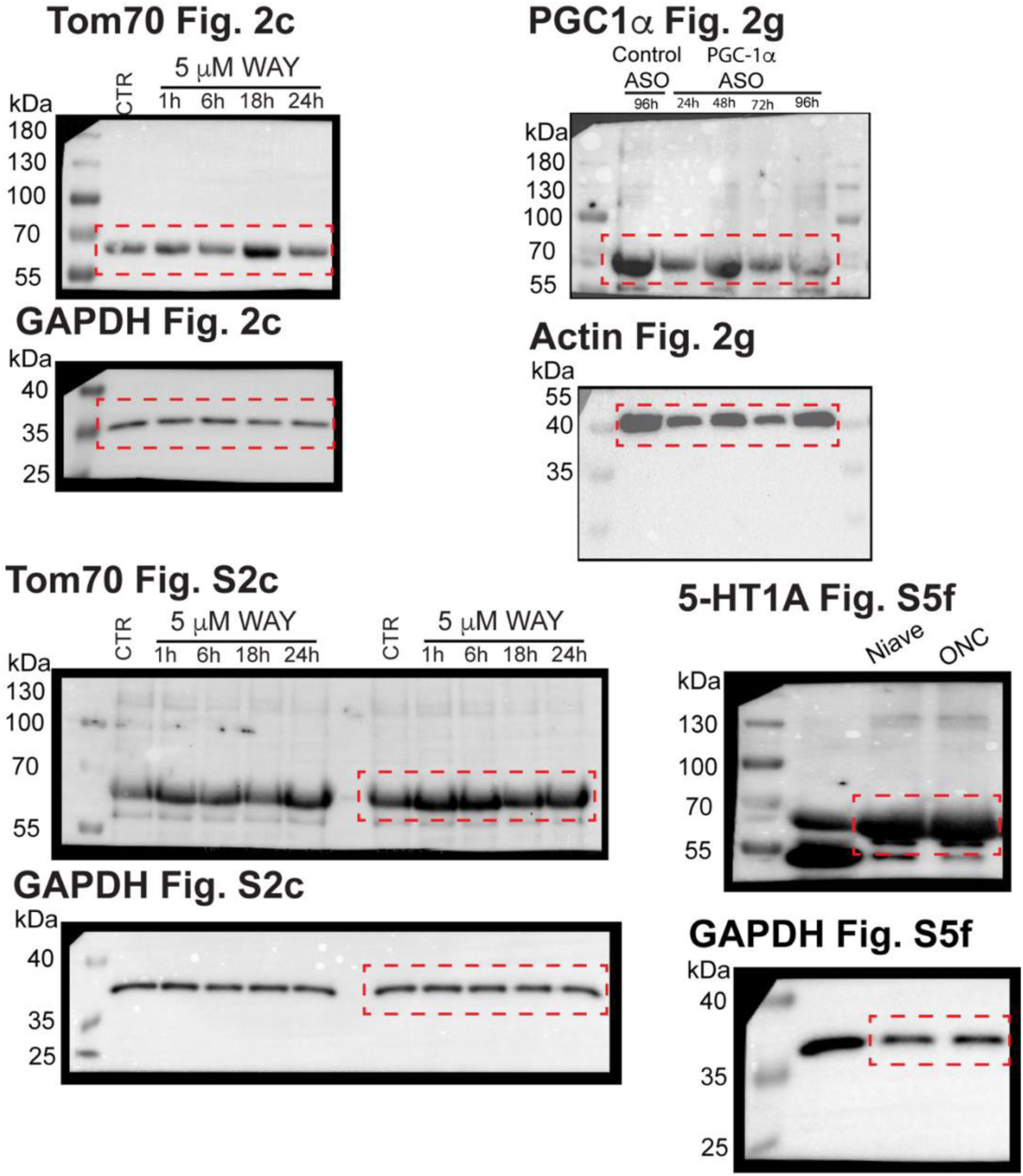
Uncropped images of representative Western blots. Uncropped Western blot membranes from corresponding blots shown in the main figures. Molecular weight markers are included for reference. Red boxes highlight the protein bands that are presented in the main figures. These full blots are provided to ensure transparency and confirm the specificity and integrity of the detected signals.

**Fig. S8.**
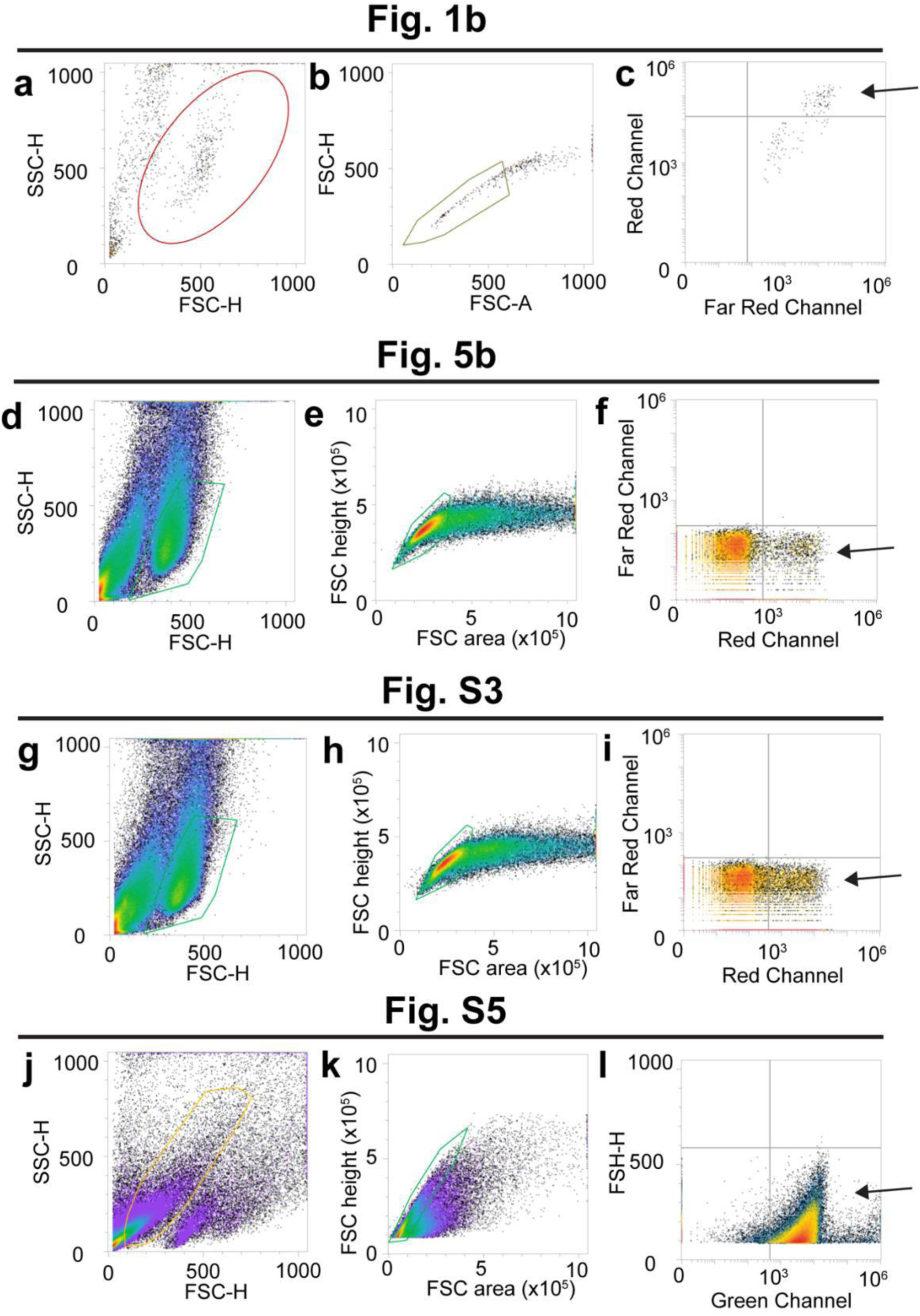
Gating strategy for flow cytometry measurements. Flow cytometry gating strategy used for analysis in the main and supplemental figures. (**a, d, g, j**) Live cells were first gated based on forward scatter (FSC) versus side scatter (SSC) profiles as demonstrated by us^22^. From the gated live cell population, (**b, e, h, k**) singlets were identified using FSC-area versus FSC-height to exclude doublets and debris. These singlet populations were further analyzed for specific parameters: (**c**) mitochondrial mass was assessed by measuring MTDR fluorescence intensity (related to Fig. 1a, b); (**f, i, l**) the percentage of cells expressing fluorescent markers such as TdTomato (**f, i,** related Fig. 5b - d; S3b, c) or immunolabelled Tuj1 (Alexa Fluor 488) (**l,** related to Fig. S5e) was quantified. Arrow indicating final cell population quantified for the study.

